# Mechanical environment afforded by engineered polymer hydrogel critically regulates survival of neural stem cells transplanted in the injured spinal cord via Piezo1-mediated mechanotransduction

**DOI:** 10.1101/2025.04.15.648586

**Authors:** Hee Hwan Park, Yurim Kim, Byeong Seong Jang, Simay Genişcan, Dong Hoon Hwang, Yeojin Seo, Seung-Ah Jee, Hyo Gyeong Seo, Hyung Soon Kim, Ariandokht Einisadr, Ho-Jeong Kim, Seolhee Lee, Sangwoo Kwon, Kyung Sook Kim, Kang In Lee, Jae Young Lee, Joo Min Park, Young-Min Kim, Soo-Chang Song, Byung Gon Kim

**Affiliations:** Ajou University School of Medicine, Department of Brain Science, Suwon, 16499, Republic of Korea; Neuroscience Graduate Program, Department of Biomedical Sciences, Ajou University Graduate School of Medicine, Suwon, 16499, Republic of Korea; Ajou University School of Medicine, Department of Neurology, Suwon, 16499, Republic of Korea; Center for Biomaterials, Korea Institute of Science and Technology, Seoul, 02792, Republic of Korea; University of Science and Technology (UST), Daejeon, Republic of Korea; Center for Cognition and Sociality, Institute for Basic Science, Daejeon, Republic of Korea; Department of Biomedical Engineering, Ulsan National Institute of Science and Technology (UNIST), Ulsan, Republic of Korea; Department of Biomedical Engineering, College of Medicine, Kyung Hee University, Seoul 130-710, Republic of Korea; ToolGen Inc., Seoul, 07789, Republic of Korea; Ajou University School of Medicine, Department of Anatomy, Suwon. 16499, Republic of Korea; Wonju Medical Industry Technovalley, Wonju 26354, Republic of Korea

## Abstract

Neural stem cell (NSC) transplantation is a promising therapeutic approach for spinal cord repair, but poor graft survival remains a critical challenge. Here, we demonstrate that the mechanical properties of the transplantation microenvironment play a crucial role in NSC survival in the injured spinal cord. While our previously engineered imidazole-poly(organophosphazene) (I-5) hydrogel effectively prevented cavity formation by promoting extracellular matrix remodeling, NSCs transplanted with 10% hydrogel exhibited poor survival. Remarkably, increasing the hydrogel concentration to 16%, which created a 5-fold stiffer matrix, significantly enhanced NSC graft survival and synaptic integration. Using *in vitro* models with controlled substrate stiffness, we found that NSCs on stiffer substrates displayed enhanced adhesion, complex morphology, and increased viability. Importantly, we identified the mechanosensitive ion channel Piezo1 as the key molecular mediator of these stiffness-dependent behaviors. CRISPR/Cas9-mediated *Piezo1* gene editing in NSCs significantly reduced graft survival *in vivo* when transplanted with 16% hydrogel, confirming that Piezo1-mediated mechanotransduction is essential for NSC survival in the injured spinal cord. Our findings reveal a previously unrecognized mechanism governing graft survival in the injured spinal cord and suggest that optimizing the mechanical properties of biomaterial scaffolds or targeting Piezo1-dependent mechanotransduction could substantially improve outcomes of cell-based therapies for neurological disorders.

## Introduction

Cell-based therapy holds significant promise for treating spinal cord injury (SCI), a devastating condition that often leads to permanent neurological deficits (*1, 2*). Several mechanisms facilitate the repair of the injured spinal cord, providing neuroprotection by modulating inflammatory responses, altering the regeneration-inhibiting environment at the injury site, and promoting remyelination in the spared white matter (*1*). In particular, neural stem or progenitor cells (NSCs) transplanted into the injured spinal cord can potentially differentiate into neurons and glial cells, replace lost tissue, provide neurotrophic support, and may help rebuild neural connections to create a new relay circuit (*3, 4*). However, a significant challenge that limits the efficacy of NSC transplantation is poor graft survival, with the majority of transplanted cells dying within days after transplantation (*5–7*). The issue of grafted cell survival is not limited to NSCs or cell transplantation for SCI; similar survival constraints have been observed with other cell types, including Schwann cells (*8*) and mesenchymal stem cells (*9, 10*), as well as in the context of transplantation for other neurological disorders (*11, 12*).

Several factors contribute to the hostile microenvironment in injured spinal cord tissue that compromises graft survival, including inflammation, oxidative stress, and lack of appropriate extracellular matrix (ECM) support (*7, 13*). To address these challenges, biomaterial scaffolds have been developed to provide structural support and a protective environment for transplanted cells (*14, 15*). Among these, hydrogels have emerged as particularly promising biomaterials due to their tunable properties, injectability, and ability to create a permissive environment for cell survival and integration (*16–18*). The physical properties of hydrogels, particularly their mechanical stiffness, significantly influence cellular behaviors such as adhesion, migration, proliferation, and differentiation (*19–21*). However, the impact of hydrogel stiffness on NSC survival after transplantation into the injured spinal cord has not been explored.

We previously engineered a polymer hydrogel composed of imidazole-poly(organophosphazene) (I-5 hydrogel) and demonstrated that the imidazole group enabled the hydrogel to minimize cavity formation by promoting ECM remodeling after SCI (*18*). While this I-5 hydrogel effectively created proteinaceous matrices at the lesion epicenter, we observed that NSCs transplanted in combination with this hydrogel still exhibited poor survival. This unexpected finding led us to investigate whether modulating the mechanical properties of the hydrogel could enhance NSC survival after transplantation. In this study, we hypothesized that increasing the stiffness of the I-5 hydrogel would improve the survival of transplanted NSCs in the injured spinal cord. Furthermore, we sought to identify the cellular and molecular mechanisms underlying the stiffness-dependent survival of NSCs, focusing on the potential role of Piezo1-mediated mechanotransduction. Our findings demonstrate a crucial relationship between mechanical cues, Piezo1 activation, and NSC survival, offering new insights for enhancing cell-based therapies for SCI and other neurological disorders.

## Results

### Transplantation of NSCs complexed with I-5 hydrogel does not improve graft survival

Our previous study reported that NSCs transplanted into the spinal cord after a contusion injury exhibit poor survival (*6*). Consistent with this finding, the majority of NSCs grafted one week after the contusion injury disappeared within 4 weeks, leaving only a few surviving cells along the cavity walls (Fig. 1A). At the lesion epicenter, we consistently observed the formation of cystic cavities of varying sizes and geometries, suggesting that the absence of ECM at the core of the lesion may contribute to their poor survival. We predicted that when NSCs are combined with I-5 hydrogel, which has been shown to prevent cavity formation by remodeling the ECM (*18*), the chances of NSC survival would improve because the hydrogel can create a protein-based matrix that allows NSCs to adhere to and interact with their surrounding matrix environment. As expected, transplantation of NSCs combined with I-5 hydrogel resulted in the formation of eosin-positive matrices at the lesion core, where in the absence of the hydrogel, we would have seen tissue defects accompanied by cystic spaces. However, contrary to our expectations, there were very few NSCs (see Fig. 1A, case #1) or only a small number of NSCs present in the newly created ECM (see Fig. 1A, case #2). Our previous study indicated that significant loss of grafted NSCs occurs as early as 3 days post-transplantation (*6*). We noticed that the number of surviving NSCs had already decreased 1 day post-transplantation (dpt), and the NSC survival was further reduced between 1- and 3- dpts (Fig. 1B). However, there was no appreciable difference in the survival between 3- and 7-day time points, suggesting that NSCs undergo substantial death within a few dpts, when I- 5 hydrogel has not yet fully degraded and active ECM remodeling is still ongoing (*18*). One possible mechanism mediating the death of grafted NSCs in the injured spinal cord involves reactive nitrogen species (*6, 22*). Nitrotyrosine antibodies were used to visualize the nitration of tissue proteins by reactive peroxynitrite molecules (*23*). At 3 dpt, we frequently observed that the GFP-positive NSCs colocalized with nitrotyrosine immunoreactivity. However, by 7 days, the proportion of nitrotyrosine-positive NSCs appeared to decrease, indicating that the NSCs were primarily exposed to reactive nitrogen species during the first few days following transplantation.

**Figure 1.**
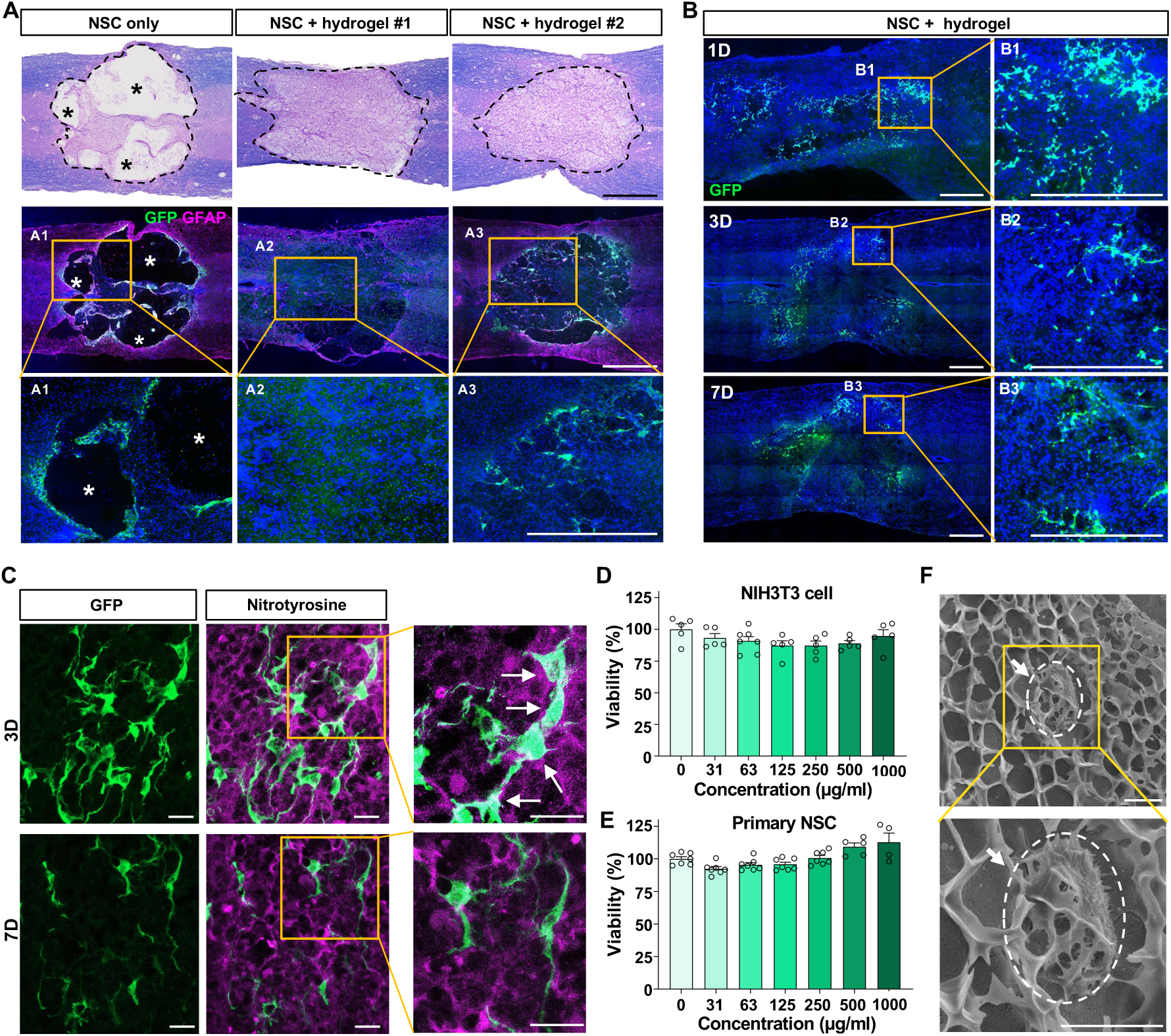
Transplantation of NSCs with hydrogel does not improve graft survival. **(A**) Representative images of longitudinal spinal cord sections obtained from animals transplanted with NSC grafts with or without hydrogel at 4 weeks after transplantation (5 weeks after injury). GFP-expressing NSCs were injected into the spinal cord at 7 days after injury. When NSCs were transplanted without hydrogel (left), the injection was done twice at 2 mm rostral and caudal to the epicenter. For transplantation with hydrogel, injection was done once at the epicenter. Huge cystic cavities were observed when NSCs were injected without hydrogel, and only very few GFP-positive NSCs were observed along the cavity walls (left). In contrast, injection of NSCs with hydrogel resulted in the formation of ECM at the lesion center. However, there was no visible GFP-positive NSCs (middle, NSC + hydrogel #1) or very few NSCs surviving within the newly created ECM (right, NSC + hydrogel #2). Boxed regions are magnified below (A1-A3). Asterisk indicates cystic cavity at the lesion epicenter. Scale bars = 500 μm. **(B)** Representative images of longitudinal spinal cord sections from animals with NSC grafts and hydrogel, taken at 1, 3, and 7 days post-transplantation (dpts). The number of surviving NSCs was already low at 1 day and further decreased at 3 and 7 days, indicating that NSC death occurs very early after transplantation. Boxed regions are magnified on the right side of the panel (B1-B3). Scale bars = 500 μm. **(C)** Representative images of surviving NSC grafts at 3 and 7 dpts. Immunostaining with anti-nitrotyrosine antibodies was employed to visualize the nitrated proteins formed by reactive peroxynitrite molecules. White arrows indicate co-localization of NSC grafts with nitrated proteins. Scale bars = 20 μm. **(D-E)** Cellular cytotoxicity assay of hydrogel solution, imidazole-poly(polyorganophosphazene) (I-5). There was no toxicity of the hydrogel at up to 1 mg/ml concentrations for both NIH3T3 cells (D) and primary NSCs (E). Error bars represent SEM. **(F)** Representative cryogenic scanning EM image of I-5 hydrogel complexed with NSCs. The size of pores formed in a gel state was smaller than that of NSCs. The white arrows indicate a NSC attached on the hydrogel. Scale bar = 5 μm.

To exclude any chance of cytotoxicity from the I-5 polymer molecules, we conducted a cell survival assay. In this assay, cells were treated with different concentrations of the I-5 polymer solution. Either the NIH3T3 fibroblast cell line or primary NSCs showed no evidence of cytotoxicity at concentrations up to 1000 μg/ml (Fig. 1D, E). It is also conceivable that the size of NSCs is smaller than the pores formed within the hydrogel, which could lead to inadequate encapsulation by the hydrogel. To investigate this possibility, a 10% I-5 hydrogel complexed with NSCs was subjected to cryogenic SEM to compare the sizes of the NSCs and the hydrogel pores (Fig. 1F). The diameter of the pores in the hydrogel ranged from one to two micrometers to approximately five micrometers, with very few pores exceeding five micrometers in diameter. In contrast, the majority of NSCs growing within the hydrogel had diameters larger than five micrometers. This indicates that grafted NSCs are unlikely to be filtered out through the hydrogel pores, reducing the likelihood that they were directly exposed to a hostile injury microenvironment. Some studies have reported that when transplanted cells pass through a syringe needle, their cellular membranes may be damaged due to shear stress, which is proportional to the viscosity of the cell suspension (*7, 24*). We speculated that NSCs complexed with viscous I-5 hydrogel might experience significant shear stress during the injection process. To investigate this, we examined whether there was an increase in the proportion of dead NSCs after the cells in the I-5 polymer complex underwent the injection procedure using the same apparatus and techniques. However, our results indicated that there was no change in the number of dead NSCs before and after the injection (Fig. S1). We also investigated whether the survival of grafted NSCs could be enhanced by incorporating growth factors into the hydrogel by activating survival pathways in NSCs. Our previous study demonstrated that IGF-1 signaling is crucial for the survival of NSC grafts (*25*). However, we did not observe significant improvement in survival when NSCs were combined with hydrogel containing IGF-1 (Fig. S2A). ECM proteins that promote cellular adhesion are commonly utilized to improve graft survival when engineered hydrogels are employed to support therapeutic cell transplantation (*24, 26*). We attempted to incorporate laminin, an ECM protein frequently used to enhance cell attachment, but we failed to increase graft survival (Fig. S2B).

### Concentration of hydrogel determines the mechanical stiffness and influences the survival of NSC grafts in the injured spinal cord

There is mounting evidence that the physical properties of biological molecules exert crucial roles in determining cellular behaviors (*27–29*). For example, matrix stiffness can modulate cancer cell metastasis, intravasation, and metabolic pathway (*30–32*). Moreover, mechanical cues influence NSC behavior based on substrate stiffness (*33–35*). Recent studies emphasize the significance of the physical stiffness of engineered substrates that utilize artificial biomaterial scaffolds (*19*). Therefore, we hypothesized that modulating the mechanical stiffness of the I-5 hydrogel may influence the survival of grafted NSCs transplanted as a complex with hydrogel. Assessment of the mechanical properties of the I-5 hydrogel using a rheometer indicated that the Young’s modulus, which measures the stiffness of solid materials, was found to be 1.6 kPa at 37°C for 10% I-5 hydrogel (Fig. 2A). This value is markedly lower than the reported value in the rat spinal cord, which is approximately 10 kPa (*36*). Hydrogel stiffness can be controlled by adjusting the polymer concentration (*37–39*). Increasing the hydrogel concentration from 10% to 16% resulted in the stiffness of the 16% hydrogel being approximately 5 times greater than that of the 10%, reaching 9.3 kPa at 37°C (Fig. 2A). We found that transplanting NSCs complexed with 16% hydrogel led to a notably improved survival of grafted NSCs compared to transplantation with 10% hydrogel (Fig. 2B). Large clusters of GFP-positive NSC grafts were frequently located along the periphery of the hydrogel-created ECM (see Fig. 2B, case #1). In some instances, nearly the entire area of the ECM was occupied by NSC grafts (see Fig. 2B, case #2). In 11 animals with NSC transplants complexed with 16% hydrogel, 8 (72.7%) animals displayed signs of surviving grafts (Fig. 2B). In contrast, transplantation was considered successful in only 3 out of the 8 animals (37.5%) that received transplants with 10% hydrogel. Additionally, the volume of NSC grafts was approximately 6 times greater in the group using 16% hydrogel compared to the 10% hydrogel group (Fig. 2C).

**Figure 2.**
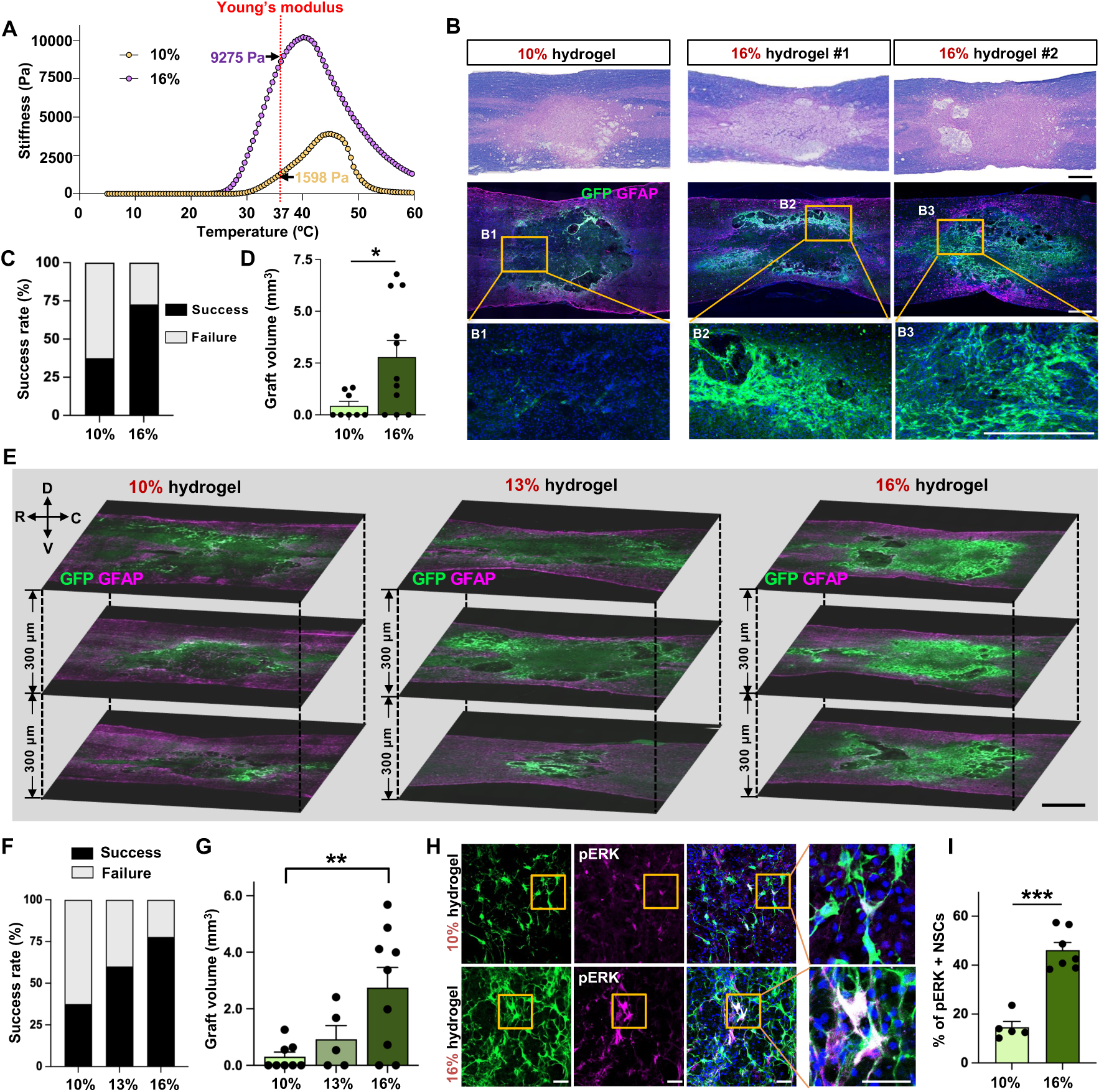
Concentration of hydrogel determines the mechanical stiffness and influences the survival of NSC grafts in the injured spinal cord. **(A)** Stiffness of 10% and 16% I-5 hydrogel was measured using a rheometer over a temperature range of 5 to 60 °C. **(B)** Spinal cord lesion areas were visualized using eriochrome cyanine and eosin staining (top panel). Surviving NSCs were identified as GFP-positive cells (green, middle panel), and lesion borders were demarcated by GFAP immunostaining (magenta, middle panel). Enhanced graft survival was noted when NSCs were transplanted with 16% hydrogel (two representative images on the right) compared to the 10% hydrogel (left). Boxed regions in the middle panel are magnified below (B1-B3). Scale bar = 500 μm. **(C, D)** Quantitative graphs comparing the rate of graft success **(C)** and the volume of GFP-positive NSC grafts **(D)**. Graft success was defined as the presence of noticeable GFP positive NSC grafts in at least one tissue section. Error bars represent SEM. * indicates *p* < 0.05 by unpaired t-test. N = 8 and 11 for 10% and 16% hydrogel groups, respectively. **(E)** Representative images of the three longitudinal spinal cord sections containing the lesion epicenter are displayed with a 300 μm interval along the dorsal-ventral (D-V) axis. R indicates rostral and C caudal direction. Animals were transplanted with GFP-positive NSCs with 10%, 13%, or 16% hydrogel after 1 week post-injury. Samples were collected 4 weeks post-transplantation (5 weeks post-injury). Surviving NSCs were identified as GFP-positive cells (green), and lesion borders were marked by GFAP immunostaining (magenta). Scale bar = 1000 μm. **(F-G)** Quantitative analysis of the graft success rate **(F)** and the volume of GFP-positive NSC grafts (G). ** indicates *p* < 0.01 by one-way ANOVA followed by Tukey’s *post hoc* analysis. Error bars represent SEM. N = 8, 5, and 9 animals for 10%, 13, and 16% hydrogel groups, respectively. **(H)** Representative images of phospho-ERK immunostaining with surviving NSC grafts. Boxed regions are magnified on the right side. Scale bar = 50 μm. **(I)** Quantitative graph showing the proportion of GFP and phospho-ERK double-positive cells. Error bars represent SEM. *** indicates *p* < 0.001 by unpaired t- test. N = 5 for 10% and N = 7 animals for 16% hydrogel group.

To confirm that a higher percentage of I-5 hydrogel supports the survival of NSCs, we designed a new experiment involving a different set of animals. In this experiment, the concentration of the hydrogel was titrated to three different levels; 10%, 13%, and 16%, and NSCs were transplanted as a complex with each of these hydrogel concentrations. The influence of hydrogel concentration on NSC survival was also demonstrated in this replication experiment as the degree of NSC graft survival was correlated with hydrogel concentration (Fig. 2E). The proportion of animals with successful transplantation increased with higher hydrogel percentages, with survival rates of 37.5% at 10%, 60.0% at 13%, and 77.8% at 16% (Fig. 2F). A one-way ANOVA revealed a significant difference in the volume of NSC grafts among the three groups with different hydrogel concentrations (*F*_(2, 19)_ = 5.96, *p* < 0.01), and *post hoc* analysis showed a significant difference between the 10% and 16% groups (Fig. 2G). We have previously demonstrated that the phosphorylation of ERK, a downstream kinase involved in growth factor-dependent prosurvival signaling, can serve as a marker for surviving NSCs following transplantation (*6*). We counted the number of phospho-ERK (pERK) positive NSCs in animals with successful transplantation and found that the proportion of pERK positive NSCs was markedly increased in animals with transplantation using 16% hydrogel compared to those using 10% hydrogel (Fig. 2H, I). Since neuroinflammation at the injury site can significantly impact the survival of transplanted NSCs (*40*), we investigated whether varying hydrogel concentrations affected the activation of inflammatory cells at the injury site. Consistent with our previous report (*18*), the intensity of immunoreactivity against IBA-1, a marker for myeloid cell lineage, was notably reduced in animals that received NSC transplantation combined with hydrogel, compared to those that underwent NSC transplantation alone (Fig. S3). However, there were no significant differences in IBA-1 immunoreactivity between the groups with different hydrogel concentrations (Fig. S3).

### Influence of hydrogel concentration on NSC integration into the host spinal cord

Previous studies have shown that the mechanical environment in hydrogel scaffolds influences differentiation of stem cells grown on them (*41, 42*). Therefore, we assessed the differentiation outcomes of NSCs transplanted with varying concentrations of hydrogels. Approximately 30% of the grafted NSCs differentiated into Tubb3-positive neuronal lineage, while a similar percentage exhibited GFAP-positive astrocytic differentiation (Fig. 3A, B). There were no significant differences between the groups with 10% and 16% hydrogels.

**Figure 3.**
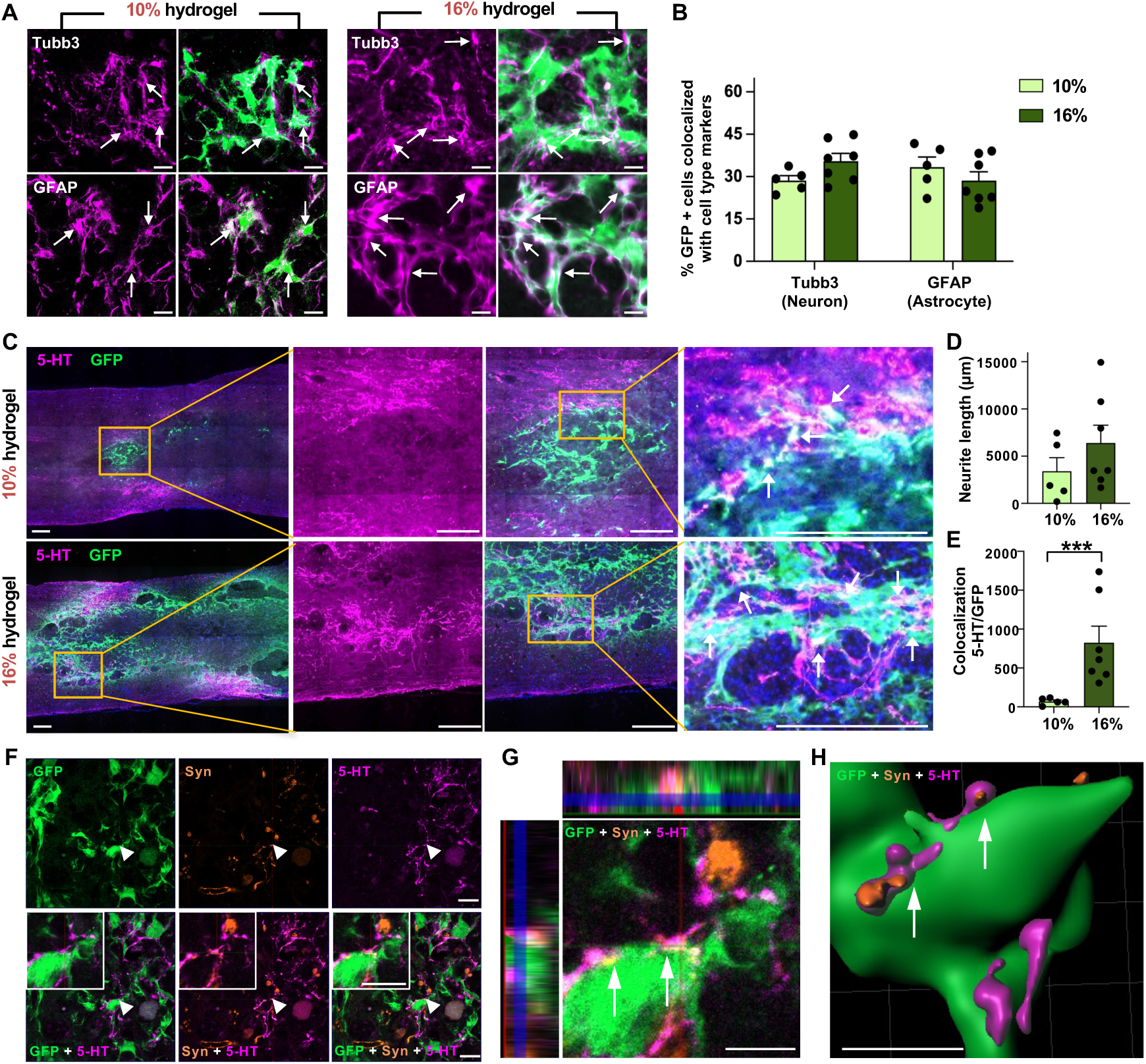
Influence of hydrogel concentration on NSC integration into the host spinal cord. **(A)** Representative images of phenotypic differentiation of NSC grafts into neuronal (Tubb3+, upper panel) and astrocytic lineage (GFAP+, lower panel). White arrows indicate co-localization of NSC grafts with each cell type marker. Scale bar = 20 μm. **(B)** Quantitative analysis of differentiation percentages in NSC grafts. N = 5 and 7 for 10% and 16% hydrogel groups, respectively. Error bars represent SEM. **(C)** Representative images of longitudinal spinal cord sections immunostained with anti-5-HT antibody, serotonergic axon (5-hydroxytryptamine) fiber marker. Boxed regions are magnified on the right side. White arrows indicate co-localization of 5-HT axons (magenta) with NSC grafts (green). Scale bars = 200 μm. **(D–E)** Quantification of 5-HT+ axon length growing within NSC grafts **(D)** and co-localization with NSC grafts **(E)**. *** indicates p < 0.001 by unpaired t-test. N = 5 and 7 animals for 10% and 16% groups, respectively. Error bars represent SEM. **(F)** Representative co-localization images of NSC grafts (GFP, green) with host 5-HT axons (5-HT, magenta) and presynaptic vesicle (synaptotagmin, orange). White arrowheads indicate tripartite co-localization Scale bars = 10 μm. **(G–H)** Orthogonal **(G)** and 3D-rendered **(H)** images showing synaptic integration of NSC grafts with 5-HT+ presynaptic axons (white arrows). Scale bar = 5 μm.

Our previous study demonstrated that the fibrotic microenvironment within the hydrogel-created ECM hinders host axons from growing into the newly assembled matrix (*43*). We evaluated the expression of fibrotic proteins by quantifying fibronectin immunoreactivity and found no significant difference between the two groups (Fig. S4A, B). In addition, there were no notable differences in the extent of Picrosirius staining, which visualizes total collagen fibrils (Fig. S4A, C). These results suggest that neither the higher percentage of hydrogel nor the enhanced survival NSC grafts significantly affected the formation of fibrotic microenvironment. When we visualized the raphespinal axons from the brainstem serotonergic neurons, the growth of 5-hydroxytryptamine (5-HT) axons into the grafts was primarily confined to the regions near the boundary between the grafts and the host spinal cord (Fig. 3C). The quantification of axon lengths growing within the GFP-positive grafts showed a trend toward an increase in animals that received NSCs in combination with 16% hydrogel. However, there was no statistically significant difference between the two groups (Fig. 3D). We also examined the interaction between the 5-HT axons growing into the grafts and GFP-positive NSCs. The number of these interactions, reflected by the colocalization of 5-HT and GFP immunoreactivities (arrows in Fig. 3C), was significantly higher in the 16% hydrogel group than the 10% group (Fig. 3E), probably due to a higher number of surviving NSCs in this group. The 5-HT axons appeared to establish synaptic contacts with the GFP-positive cells, as we observed a close apposition between the GFP+ cells and the 5-HT axon terminals, which were colocalized with a presynaptic marker, synaptotagmin (Fig. 3F-H). Collectively, these results indicate that enhance survival of NSCs via 165 hydrogel improved synaptic integration of transplanted NSCs.

### Influence of the substrate stiffness on the adhesive properties and the viability of cultured NSCs

Hydrogels with varying concentrations may affect the survival of neural stem cells (NSCs) independently of changes in mechanical stiffness. For instance, different concentrations of hydrogels could influence degradation rates or the speed of gelation (*44*), both of which are crucial for encapsulating or protecting NSCs within the hydrogel. To examine whether different hydrogel concentrations affect degradation rates *in vivo*, we subcutaneously injected 10% and 16% I-5 hydrogels mixed with a fluorescent dye into the left and right dorsal skin of mice and monitored the fluorescence signals from the hydrogel (Fig. S5). The fluorescence intensity began to decrease as early as 3 days post-injection and continued to decline until 14 days, when signals were barely detectable. However, we did not find any significant differences in the degradation kinetics between the 10% and 16% hydrogels (Fig. S5). We also compared the speed of gelation by injecting 10% and 16% I-5 hydrogels mixed with dye into a jar containing warm water at 37°C. The injected hydrogel sank to the bottom of the jar and transformed into a gel-like material within a few seconds (Movie S1). No noticeable difference in gelation speed was observed between the hydrogels of different concentrations.

The results above led us to hypothesize that the mechanical properties of the hydrogel, determined by its concentration, could significantly influence the viability of NSCs surrounded by the hydrogel. To test this hypothesis *in vitro*, we utilized a culture model in which the mechanical stiffness of the substrate on which NSCs are grown is precisely controlled (*45, 46*). NSCs were plated onto polyacrylamide (PAA) hydrogel coverslips with elastic moduli of 25, 2, 0.5, and 0.2 kPa (Fig. 4A, B). We first examined the influence of hydrogel substrates with varying levels of stiffness on the adhesion and cellular morphology of cultured NSCs. There was a significant group difference in the number of NSCs attached onto substrates with different elastic moduli (*F*_(3, 12)_ = 8.198, *p* < 0.01), with the number of NSCs attached onto the most rigid substrate (25 kPa) significantly higher than that on softer substrates (0.5 and 0.2 kPa) (Fig. 4B, C). Additionally, NSCs grown on softer substrates (0.5 and 0.2 kPa) exhibited a more round morphology with fewer cytoplasmic extensions, while NSCs grown on stiffer substrates (25 and 2 kPa) displayed a polygonal shape with more prominent processes. To quantitatively compare the differences in cellular morphology, we measured the spreading area and perimeter of adhered NSCs. Significant differences were observed between the groups for both measurements: *F*_(3, 12)_ = 22.63, *p* < 0.001 for spreading area, and *F*_(3, 12)_ = 20.88, *p* < 0.001 for perimeter. *Post hoc* analysis indicated that both the spreading area and perimeter of NSCs on stiffer substrates (25 and 2 kPa) were significantly larger than those on softer substrates (0.5 and 0.2 kPa) (Fig. 4D, E).

**Figure 4.**
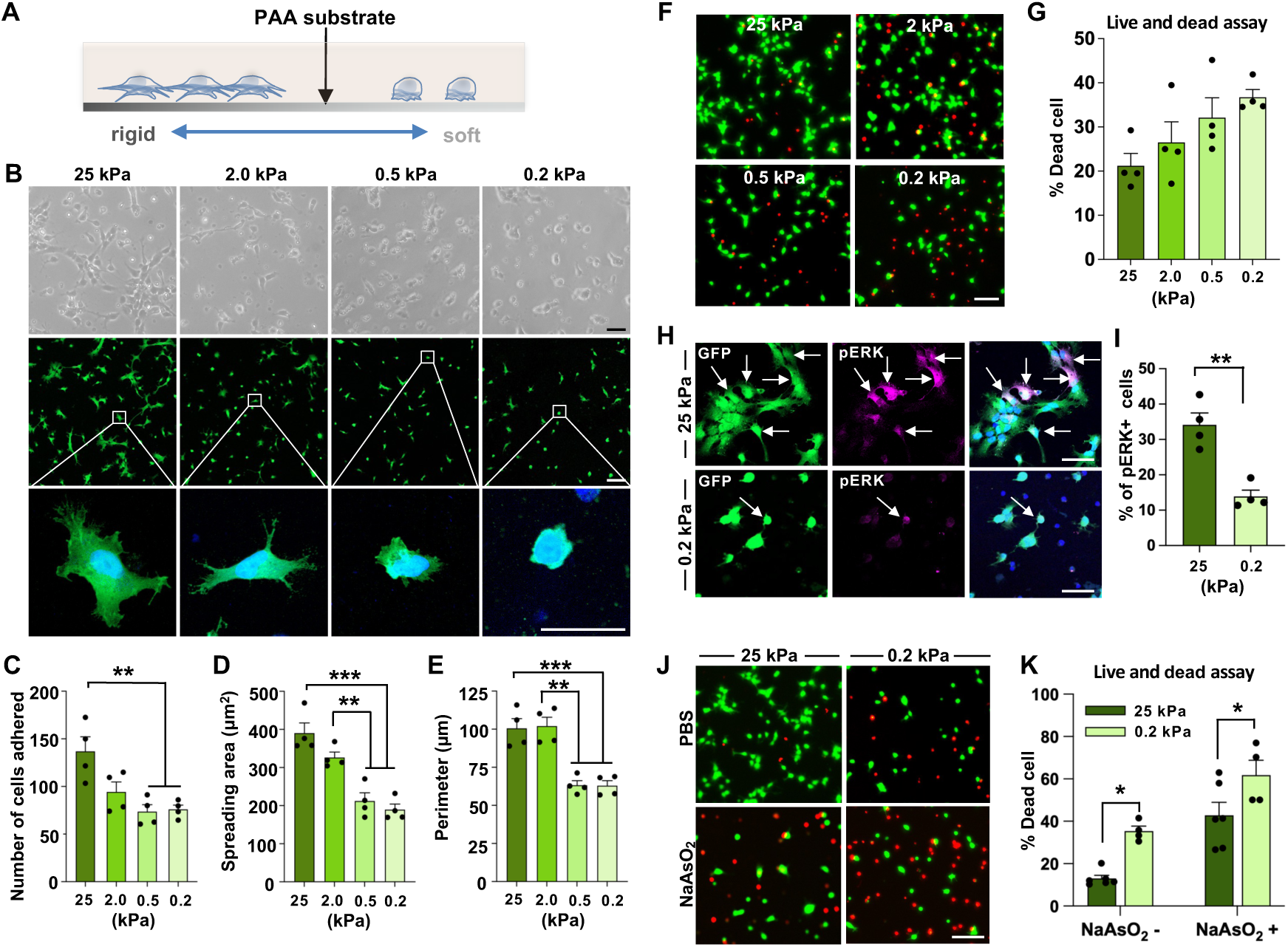
Influence of the substrate stiffness on the adhesive properties and the viability of cultured NSCs. **(A)** A diagram depicting the *in vitro* culture system providing polymer matrix with varying degree of mechanical stiffness. PAA = polyacrylamide. **(B)** Representative images of NSCs obtained during adhesion assay. NSCs expressing GFP were grown on substrates with mechanical stiffness ranging from 25 kPa to 0.2 kPa. Upper panel: Phase-contrast images. Middle panel: NSCs were visualized using GFP fluorescence. Boxed regions show magnified cell morphologies below. Scale bars = 20 μm. **(C-E)** Quantitative graphs comparing the number of cells adhered (C), areas of cell spreading (D), and the perimeter of cell boundary (E). Each dot represents an independent culture replicate, with each replicate being an average value from three coverslips. Error bars represent SEM. ** and *** indicates p < 0.05 and p < 0.001 by one-way ANOVA followed by Tukey’s *post hoc* analysis. (**F-G)** Representative images of NSCs for cell survival assay. NSCs were cultured on hydrogel substrates ranging from 25 kPa to 0.2 kPa stiffness for 24 hrs. Subsequently, cell survival assay was performed. Live cells are labeled by Calcein-AM (green), and dead cells are labeled by Ethidium Homodimer-1 (red). Quantitative graph depicting the percentage of dead cells **(G)**. Each dot represents an independent culture replicate, with each replicate being an average value from three coverslips. Error bars indicate SEM. Scale bars = 100 μm. **(H-I)** Representative images of phospho-ERK immunostaining **(H)**. Arrows indicate GFP positive cells colocalized with phospho-ERK (pERK, magenta) expression. The total number of NSCs was divided by pERK positive cells **(I)**. Error bars represent SEM. ** indicates *p* < 0.01 by unpaired t-test. Each dot represents an independent culture replicate, with each replicate being an average value from three coverslips. Scale bar = 50 μm. (**J-K)** Representative images of NSCs for cell survival assay with sodium arsenite treatment. Sodium arsenite (NaAsO_2_) was used to induce cellular stress for 24 h **(J)**. Quantitative graph of percentage of dead cells **(K)**. N = 6 and 4 for 25 kPa and 0.2 kPa groups, respectively. Error bars represent SEM. * indicates *p* < 0.05 by unpaired t-test. Error bars represent the SEM. Scale bars = 100 μm

Next, we examined how substrate stiffness affects cellular viability. Cell viability was measured using calcein AM for live cells and ethidium homodimer-1 for dead cells. We observed a trend toward an increase in the percentage of dead cells as stiffness decreased, with the highest percentage of dead cells found in the lowest stiffness condition (0.2 kPa) (Fig. 4F, G). However, the differences between groups were not statistically significant (*F*_(3, 12)_ = 3.45, *p* = 0.0514). We compared the expression of pERK, a marker of NSC viability post-transplantation, between NSCs grown on stiff (25 kPa) and soft (0.2 kPa) hydrogel substrates (Fig. 4H, I). The percentage of NSCs expressing pERK was significantly higher on the 25 kPa substrate than on the 0.2 kPa substrate, indicating more robust activation of pro-survival signals in NSCs on stiffer substrates. To assess the NSC viability in an environment that mimics injured spinal cord, we treated NSCs with sodium arsenite (NaAsO_2_) to induce cellular stress (*47*). The effects of substrate stiffness on NSC viability were significantly observed in the NaAsO_2_-treated conditions (Fig. 4J, K), similar to those without treatment (*F*_(1, 16)_ = 17.77, *p* < 0.001 by two-way ANOVA), suggesting that the mechanical environment may also play a role in supporting NSC viability under conditions of cellular stress.

### Actin polymerization determines the substrate stiffness-dependent cellular elasticity and intracellular calcium dynamics in NSCs

If the mechanical stiffness of a hydrogel substrate affects various cellular behaviors, it should also influence the mechanical properties of NSCs (*48*). We compared the cellular elasticity of NSCs grown on hydrogel substrates with varying mechanical stiffness using atomic force microscopy (AFM) (*49*) (Fig. S6A). We first measured the elastic modulus (EM) of the PAA hydrogel substrates. The AFM used for this study could measure the EM of the 12kPa substrate and higher, but not detect any measurable elasticity of the substrates softer than 12kPa. The measured EM values for the substrates with stiffness levels of 100, 25, and 12 kPa, as designated by the manufacturer, closely aligned with those provided by the manufacturer (Fig. S6B). NSCs cultured on a substrate with 100 kPa exhibited approximately 20 kPa elasticity (Fig. 5A), and the cellular elasticity is decreased proportionally on softer substrates. Since filamentous actin (F-actin) is a major determinant of cellular elasticity (*50*), we visualized the F-actin using the phalloidin conjuaged with a fluorescent dye. The F-actin signals were notably intense along the NSC membrane on the 25 kPa substrate (Fig. 5B). In these cells, F-actin fibers appeared as linearly running bundles, reminiscent of stress fibers, within the cytoplasmic compartment. In contrast, discernible F-actin fibers were barely detectable in the submembranous or intracellular compartments in NSCs on the 0.2 kPa substrate, resulting in a significant reduction of the F-actin signal intensity (Fig. 5B, C). To test if actin polymerization plays a role in the stiffness-dependent elasticity, NSCs on either 100 kPa or 12 kPa substrates were treated with the actin depolymerizing agent, cytochalasin D (cytoD). CytoD effectively disrupted the F-actin fiber network in NSCs in a dose-dependent manner (Fig. S7). The cellular elasticity of NSCs cultured on the 100 kPa substrate was reduced to nearly half following cytoD treatment (Fig. 5D), and CytoD treatment in the 12 kPa condition also significantly decreased cellular elasticity. However, the extent of the drug’s effect was less pronounced compared to NSCs grown on the stiffer substrate. A two-way ANOVA analysis revealed that both the drug treatment and substrate stiffness had statistically significant effects on the EM of NSCs (drug treatment, *F*_(1, 86)_ = 52.1, *p* < 0.001; stiffness, *F*_(1, 86)_ = 111.5, *p* < 0.001). Furthermore, the interaction between the two factors was also statistically significant (*F*_(1, 86)_ = 10.0, *p* < 0.01), suggesting that actin depolymerization had a greater effect on cellular elasticity in NSCs grown on stiffer substrates.

**Figure 5.**
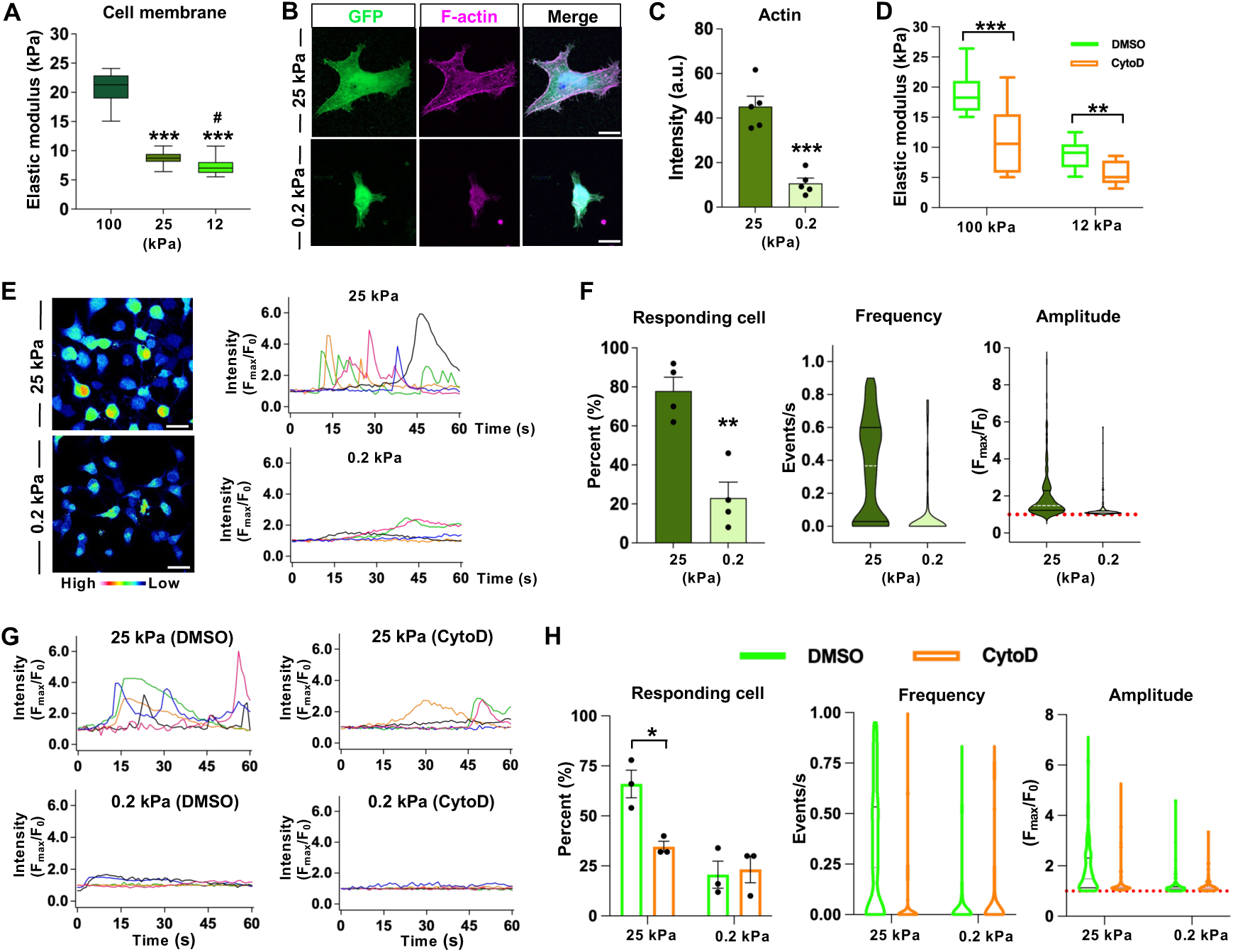
Actin polymerization determines the stiffness-dependent cellular elasticity and intracellular calcium dynamics in NSCs. **(A)** Rheological analysis of the elastic modulus of NSCs cultured on hydrogel substrates with stiffness of 100, 25, and 12 kPa. Atomic force microscopy (AFM) was used to measure cellular membrane stiffness. *** indicates *p* < 0.001 compared to 100 kPa, and # indicates *p* < 0.05 compared to 25 kPa group by one-way ANOVA followed by Tukey’s *post hoc* analysis. N= 20, 25, and 20 cells for 100 kPa, 25 kPa and 12 kPa, respectively. **(B-C)** Representative images of filamentous actin (F-actin) visualization in NSCs (B). F-actin was stained using Alexa Fluor 594-Phalloidin. Quantification graph of actin staining intensity (C). Each dot represents an independent culture. *** indicates *p* < 0.001 by unpaired t-test. Scale bar = 5 μm. **(D)** Changes in NSC plasma membrane elastic modulus measured by AFM following treatment with 20 μM of cytochalasine D (CytoD), actin destabilizer. N = 20 and 30 cells for 100 kPa and 12 kPa (DMSO) groups, and N = 20 for both 100 kPa and 12 kPa (CytoD) group. *** indicates *p* < 0.001 and ** indicates *p* < 0.05 by two-way ANOVA followed by Bonferroni’s *post hoc* analysis. **(E)** Intracellular calcium oscillations in NSCs. Left: representative calcium intensity maps (red: high level of Ca2+, blue: low level of Ca2+). Right: representative plots of calcium oscillations. Each color represents an individual cell. **(F)** Quantitative graphs of intracellular calcium oscillation parameters: responding cells (%), frequency (events/sec), and amplitude (F_max_/F_0_). ** indicates *p* < 0.01 by unpaired t-test. N = 200 and 190 cells from 4 independent cultures for 0.25 kPa and 25 kPa groups. Each dot represents an independent culture. **(G)** Intracellular calcium oscillation after cytochalasin D treatment. **(H)** Quantification graphs of intracellular calcium oscillation parameters: responding cells (%), frequency (events/sec), and amplitude (F_max_/F_0_). * indicates *p* < 0.05 by unpaired t-test. N = 150 cells for 0.2 kPa + DMSO; N = 130 cells for 0.2kPa + cytoD; N = 150 cells for 25 kPa + DMSO; and N = 150 cells for 25 kPa + cytoD group. Each dot represents an average value from an independent culture.

The mechanical environment can modulate intracellular calcium ion (Ca^2+^) oscillations (*51-53*). In addition, the dynamics of the actin cytoskeleton are closely linked to these Ca^2+^ oscillations (*54, 55*). Therefore, we examined whether the intracellular Ca^2+^ oscillations in NSCs are differentially regulated depending on the stiffness levels of a hydrogel substrate on which NSCs are grown. Many NSCs grown on a plastic dish exhibited spontaneous Ca^2+^ spikes, visualized by Fluo-4 AM, lasting for several seconds, and these spikes frequently oscillated during the 1-min imaging session (Movie S2). These spontaneous Ca^2+^ oscillations were also observed to a similar extent in NSCs grown on the 25 kPa substrate (Fig. 5E, Movie S3), with almost 80% of NSCs showing at least one transient Ca^2+^ wave. In contrast, the Ca^2+^ activities were largely attenuated in NSCs on 0.2 kPa substrate with a significantly lower number of Ca^2+^ spikes onbserved in the 0.2 kPa condition (Fig. 5F). The frequency and the amplitude of observed Ca^2+^ spikes were also markedly lowered in NSCs in the 0.25 kPa than those in the 25 kPa condition (25 kPa vs 0.25 kPa; frequency, 0.36/s ± 0.29 vs 0.08/s ± 0.19; amplitude, 0.98 ± 1.28 vs 0.25 ± 0.51). Next, we determined whether the mechanical stiffness-dependent intracellular Ca^2+^ oscillations are mediated by actin polymerization. CytoD treatment markedly attenuated the oscillatory Ca^2+^ activities in NSCs grown on the 25 kPa substrate (Movies S4), while NSCs on the 0.25 kPa substrate did not show considerable changes in response to cytoD treatment (Fig. 5G). A two-way ANOVA analysis revealed that both the drug treatment and substrate stiffness had statistically significant effects on the number of responding NSCs (drug treatment, *F*_(1, 8)_ = 5.654, *p* < 0.05; stiffness, *F*_(1, 8)_ = 22.09, *p* < 0.01). The interaction between the two factors was statistically significant (*F*_(1, 8)_ = 7.954, *p* < 0.05), indicating that cytoD treatment significantly influenced Ca^2+^ oscillation activity in NSCs grown on stiffer substrates (Fig. 5H). In the 25 kPa condition, the frequency and amplitude of Ca^2+^ spikes were substantially reduced by cytoD in NSCs (DMSO vehicle vs cytoD; frequency, 0.30/s ± 0.31 vs 0.11/s ± 0.23; amplitude, 1.84 ± 1.01 vs 1.48 ± 0.80). In contrast, cytoD treatment did not produce significant changes in the 0.25 kPa condition.

### Involvement of *Piezo1* in the stiffness-dependent adhesive behavior of cultured NSCs

Mechanosensitive ion channels translate changes in a mechanical environment into ionic flow, responding to extracellular physical cues and influencing various fundamental physiological processes at both the organismal and cellular levels (*56*). We screened the expression of *Piezo1* and various transient receptor potential (TRP) channels in NSCs, which are mechanosensitive ion channels known to be expressed in mammalian stem cells (*57*). Substantial mRNA expressions of *Piezo1*, *TRPC1*, and *TRPP2*, but not *TRPV4* and *TRPA1*, in NSCs were observed compared to the rat brain and lung tissues as positive controls (Fig. 6A). Among these, *TRPP2* had the highest expression level in NSCs, followed by *TRPC1* and *Piezo1* (Fig. 6B). To examine the functional implications of these mechanosensitive ion channels in the adhesive properties of cultured NSCs (see Fig. 4B-E), NSCs grown on a stiff 25 kPa substrate were treated with various inhibitors targeting these mechanosensitive channels. First, amiloride hydrochloride, a TRPP2 channel blocker, did not significantly impact the number of adhered NSCs at a concentration of up to 1 μM, but 10 μM of amiloride hydrochloride slightly decreased the number of adhered cells. A significant group difference was observed (*F*_(3, 8)_ = 5.850, *p* < 0.05 by one-way ANOVA), and *post hoc* analysis showed a significant decrease by amiloride hydrochloride only at 10 μM concentration (Fig. S8A-D). However, the spreading area and perimeter, which are parameters of cultured NSC morphology, were not affected (Fig. S8C, D). Treatment with Pico145, a specific inhibitor of TRPC1/4/5, did not influence the number or morphology of adhered NSCs, even at the highest concentration (Fig. S8E-H), ruling out the potential involvement of TRPC1. In contrast, when GsMTx4, an inhibitor of mechanosensitive ion channels belonging to the Piezo and TRP families was treated, the polygonal shape of NSCs, characterized by notable cellular branches, became rounder with fewer processes in response to higher concentrations of GsMTx4 (Fig. 6C), similar to NSCs on a softer substrate (see Fig. 4B). A significant group difference in the number of adhered NSCs was observed between different concentrations of GsMTx4 treatment (*F*_(3, 8)_ = 13.60, *p* < 0.01), and *post hoc* analysis showed significant decreases by GsMTx4 at 0.5 and 2.5 μM concentrations (Fig. 6D). There were significant differences also in both the spreading area and perimeter among the groups: *F*_(3, 8)_ = 16.26, *p* < 0.001 for spreading area, and *F_(_*_3, 8)_ = 6.189, *p* < 0.05 for perimeter. *Post hoc* analysis revealed a similar result to that from the cell number analysis (Fig. 6E, F). Thus, we concluded that Piezo1 was the primary mechanosensitive ion channel responsible for the adhesive properties of NSCs on a stiff substrate among those that were substantially expressed in NSCs. To verify the functionality of the Piezo1 channel in NSCs, we performed a calcium imaging experiment after treating NSCs grown on a 0.2 kPa substrate with the Piezo1 agonist Yoda1. The treatment with Yoda1 resulted in significant and sustained Ca^2+^ influxes (Fig. 6G and Movie S5), demonstrating that the Piezo1 channel is indeed functional in NSCs.

**Figure 6.**
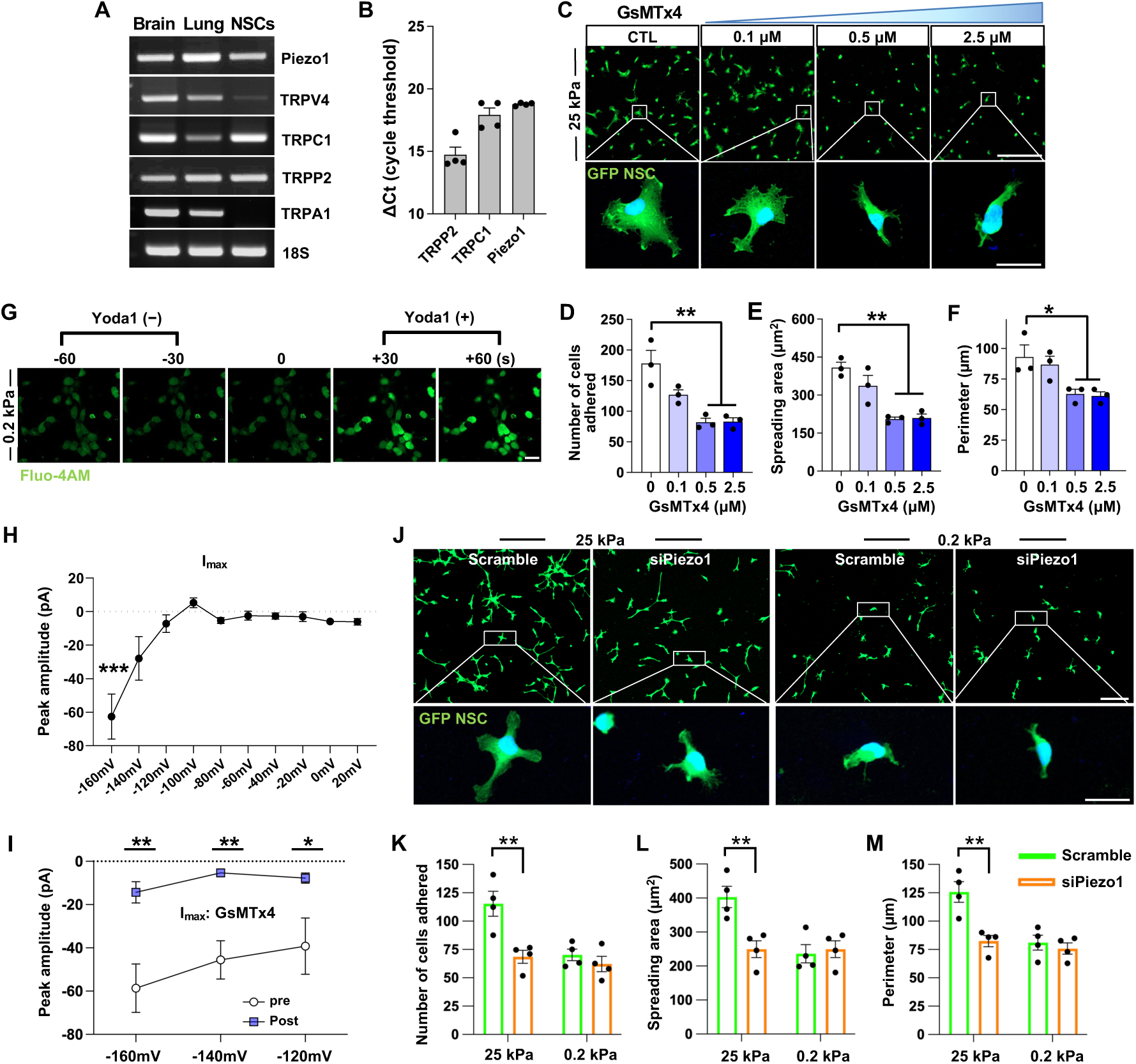
Involvement of Piezo1 in the stiffness-dependent adhesive behavior of cultured NSCs. **(A)** Representative mRNA expression of putative mechanosensitive ion channels in NSCs. Brain and lung tissues were used as positive controls. **(B)** Quantitative measurement of mRNA expression by qRT-PCR. Each dot represents an independent biological replicate. Error bars represent SEM. **(C-F)** NSC adhesion assay with GsMTx4. NSCs were treated with GsMTx4 for 24 h on 25 kPa hydrogel substrate (C) Quantitative graphs comparing the number of cells adhered (D), areas of cell spreading (E), and the perimeter of cell boundary (F). Error bars represent SEM. ** indicates *p* < 0.01 and * indicates *p* < 0.05 by one-way ANOVA followed by Tukey’s *post hoc* analysis. Three biological replicates were performed. Boxed regions are magnified below. Scale bar = 20 μm. **(G)** Representative images of Piezo1 functional assay in NSCs treated with the Piezo1 agonist (Yoda1). NSCs were grown on 0.2 kPa hydrogel substrates, and intracellular calcium uptake was visualized using Fluo4-AM. Scale bar = 20 μm. **(H)** Quantification of the peak amplitude of the inward currents (I_max_) at varying holding potentials. *** indicate *p* < 0.001 compared to the I_max_ at 0 mV holding potential by one way ANOVA followed by Tukey’s *post hoc* analysis. Each data point represents the average amplitude from 10 independent measurements using 10 cells (N = 10). **(I)** Comparison of I_max_ before and after treatment with GsMTx4, at a concentration of 5 μM. I_max_ values were obtained before drug treatment and 20 min after drug treatment from 7 independent cells (N = 7). * and ** indicate *p* < 0.05 and *p* < 0.01 by two-way ANOVA followed by *post hoc* Bonferroni test. **(J-M)** NSC adhesion assay with Piezo1 knockdown by electroporation. Scrambled siRNA was used as a control (J). Quantitative graphs comparing the number of cells adhered (K), areas of cell spreading (L), and the perimeter of cell boundary (M). Boxed regions are magnified below. ** indicates *p* < 0.01 by two-way ANOVA followed by by *post hoc* Bonferroni test. Each dot represents an independent culture replicate, with each replicate being a measurement from one coverslip. Error bars represent SEM. Scale bar = 20 μm.

To provide direct evidence that mechanical stimulation of NSCs induces ionic currents through mechanosensitive channels, we conducted whole-cell patch-clamp recordings on cultured NSCs. We activated the mechanosensitive ion channels by applying short air pulses at 7 psi positive pressure, lasting 10 ms, using a Picospritzer device and measured inward current using a voltage-clamp mode (Fig. S9). Notably, inward deflection currents in response to the pressure stimulation began to appear at −120 mV holding potentials and became more robust at higher holding potentials. The mean peak amplitude of the inward currents, I_max_, at −160 mV holding potential was approximately 60 pA and significantly larger than that at 0 holding potential (Fig. 6H). To further confirm that the observed inward currents were mediated by the Piezo1 channel, we treated NSCs with a Piezo1 blocker, GsMTx4, and compared the I_max_ values before and after the treatment. The treatment with GsMTx4 significantly reduced the magnitude of I_max_ at −120, −140, and −160 mV holding potentials (Fig. 6I), demonstrating that the pressure-induced currents were dependent on the activity of the Piezo1 channel.

To examine the role of Piezo1 in the stiffness-dependent adhesive properties of cultured NSCs, Piezo1 expression was suppressed using siRNA. The levels of Piezo1 expression in NSCs were comparable regardless of whether they were cultured on a standard plastic substrate or on PAA substrates with stiffness values of 25 kPa and 0.2 kPa (Fig. S10A). When siRNA targeting Piezo1 was treated to NSCs grown on a 25 kPa substrate, the NSCs underwent a significant morphological change from a polygonal shape to a more rounded appearance, which included a loss of cytoplasmic processes (Fig. 6J). The morphological changes were accompanied by a decrease in the number of adhered cells in the siPiezo group. In contrast, the morphology and number of NSCs cultured on a 0.2 kPa substrate showed no significant changes following Piezo1 knockdown. A two-way ANOVA analysis revealed that both the siPiezo1 treatment and substrate stiffness had statistically significant effects on the number of adhered NSCs (siPiezo1, *F*_(1, 12)_ = 13.32, *p* < 0.01; stiffness, *F*_(1, 12)_ = 11.71, *p* < 0.01) (Fig. 6K). Furthermore, the interaction between the two factors was also statistically significant (*F*_(1, 12)_ = 6.604, *p* < 0.05), indicating that Piezo1 knockdown had a more significant effect on NSCs grown on a stiffer substrate. Statistical analyses on the spreading area and the perimeter showed similar results (Fig. 6L, M), revealing the significant influence of siPiezo1 on the morphology of NSCs on a stiffer substrate. These results indicate that Piezo1 regulates the adhesive properties of NSCs when exposed to a stiffer mechanical environment.

### CRISPR/Cas9-mediated *Piezo1* gene editing abolishes the stiffness-dependent survival of NSC grafts in the injured spinal cord

The above findings led us to hypothesize that the improved survival of NSCs transplanted in a complex with higher percentage I-5 hydrogel may be attributed to the activation of Piezo1 channels in a stiffer mechanical environment. Since we found that siRNA-mediated knockdown of *Piezo1* did not last longer than a couple of days (Fig. S10B), we employed CRISPR/Cas9 gene editing for the long-term suppression of *Piezo1* in primary rat NSCs. To target rat *Piezo1*, we searched for protospacer adjacent motif (PAM) sequences (NGG for *Streptococcus pyogenes* Cas9) within the protein coding sequences of rat *Piezo1* and designed 13 different single guide RNAs (sgRNAs) to target this region (Table S1).The activities of these sgRNAs were screened after transfection of sgRNA-Cas9 RNP complexes into a rat C6 glioma cell line and selected six sequences that demonstrated an indel frequency greater than 90% using targeted deep sequencing (Fig. S11A). We then performed RT-qPCR and selected the sgRNA sequence #3 in exon1 of the *Piezo1* (lead sgRNA labeled as sgPiezo1 hereafter) based on the knockdown efficiency (Fig. S11B and Fig. 7A). Using this sgRNA, we achieved near 100% indel frequency (with most frequent edits are >38 bp deletions; Table S2) and more than 50% reduction of *Piezo1* mRNA in primary NSCs (Fig. 7B).

**Figure 7.**
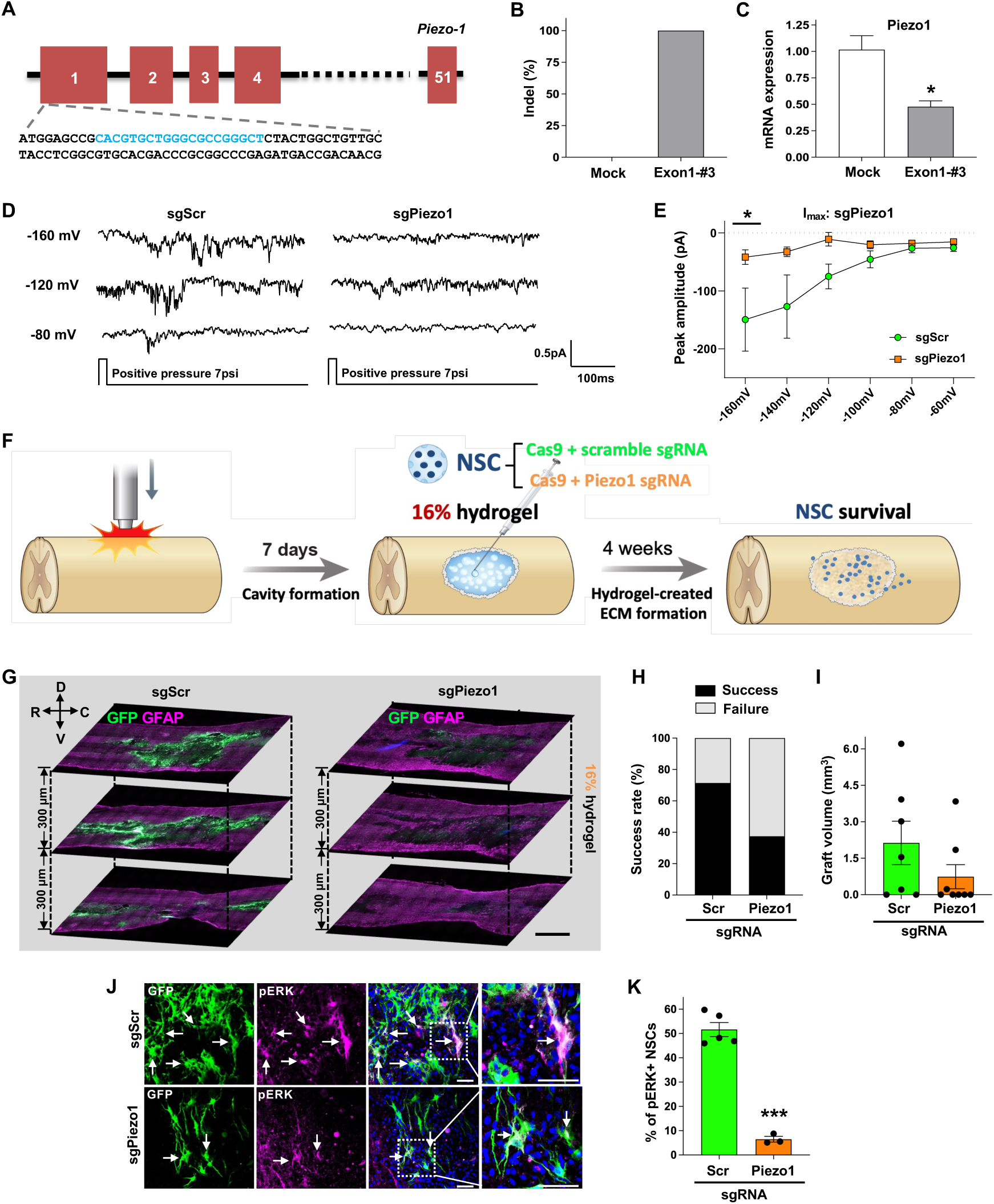
CRISPR/Cas9-mediated *Piezo1* gene editing abolishes the stiffness-dependent survival of NSCs in the injured spinal cord. **(A)** DNA sequences (shown in blue) in the proximal exon1 of the Piezo1 gene targeted by the chosen sgRNA. **(B)** Indel frequency of Crispr/Cas9 gene editing using the chosen sgRNA sequence (exon1-#3). **(C)** Measurement of Piezo1 mRNA levels by quantitative RT-PCR. * indicates p < 0.05 by unpaired t-test (N = 3 replicates per group). Error bars represent SEM. **(D)** Representative raw current traces from NSCs with control scrambled sgRNA (sgScr) and sgPiezo1 at varying holding potentials. **(E)** Quantification of the peak amplitude of the inward currents (Imax) at varying holding potentials. * indicates p < 0.05 by two-way ANOVA followed by Bonferroni’s *post hoc* analysis. N = 5 cells per group. **(F)** Schematic diagram illustrating the experimental design for Crispr/Cas9-mediated long-term Piezo1 knockdown *in vivo* study. **(G)** Representative images of longitudinal spinal cord sections obtained from animals transplanted with NSCs edited by Crispr/Cas9 (sgPiezo1) or control (sgScr). Three longitudinal spinal cord sections containing the lesion epicenter are displayed with a 300 μm interval along the dorsal-ventral (D-V) axis. R indicates rostral and C caudal direction. GFP indicates surviving NSCs (green) and GFAP (magenta) demarcates lesion areas. Scale bar = 1000 μm. **(H-I)** Quantitative graphs comparing the rate of graft success **(H)** and the volume of GFP-positive NSC grafts **(I)**. Graft success was defined as the presence of noticeable GFP positive NSC grafts in at least one tissue section. N = 7 and 8 animals for sgScr and sgPiezo1 group, respectively. Error bars represent SEM. **(J)** Representative images of phospho-ERK immunostaining with surviving NSC grafts. Boxed regions are magnified on the right side. Scale bar = 50 μm. **(K)** Quantitative graph showing the proportion of GFP and phospho-ERK double-positive cells. *** indicate *p* < 0.001 by unpaired t-test. Error bars represent SEM. N = 5 and 3 animals for sgScr and sgPiezo1.

The functional suppression of the Piezo1 channel was assessed by visualizing Ca^2+^ influx induced by Yoda1. Treatment with Yoda1 resulted in a significant increase in Ca^2+^ influx in NSCs grown on 0.2 kPa substrate that were transfected with either Cas9 alone or Cas9 in conjunction with a control scrambled sgRNA (sgScr) (Fig. S12A). However, the Ca^2+^ influx induced by Yoda1 was noticeably reduced in NSCs that were transfected with both Cas9 and sg*Piezo1* (Fig. S12A-C). To investigate whether CRISPR/Cas9-mediated editing of the *Piezo1* gene effectively inhibited ionic currents triggered by mechanical stimulation, we measured inward ionic currents in response to pressure stimulation in *Piezo1*-edited NSCs. In NSCs with control sgScr, we observed distinctive inward currents at holding potentials of - 120 mV and −160 mV (Fig. 7D), but these currents almost disappeared in NSCs with sgPiezo1. Quantitative analysis showed a significant reduction in the I_max_ at the −160 mV holding potential in NSCs with sgP*iezo1* (Fig. 7E), indicating the *Piezo1* gene editing successfully suppressed functional mechanosensitive channels.

To determine whether Piezo1 plays a crucial role in enhancing graft survival when NSCs are injected in combination with 16% I-5 hydrogel (see Fig. 2B, E), we conducted an *in vivo* experiment in which NSCs transfected with sgScr or sgPiezo1 were transplanted as a complex with the 16% I-5 hydrogel one week after an initial injury (Fig. 7F). A total of 15 animals were generated with 7 animals for sgScr and 8 animals for sgPiezo1, respectively, and these animals underwent survival analysis four weeks after transplantation. NSC grafts were successfully observed in 5 out of 7 animals (71.4%) with control sgScr, and the majority of animals with graft success showed quite a sizable engraftment (Fig. 7G-I). However, the success rate was only 37.5% (3 out of 8 animals) in sg*Piezo1* group, and the average graft volume tended to be smaller in this group (Fig. 7G-I). We counted the number of pERK positive NSCs in animals with successful transplantation and discovered that the proportion of pERK positive NSCs was significantly reduced in animals with Piezo1 gene editing (Fig. 7J, K).

## Discussion

This study was motivated by the observation that transplanting NSCs complexed with 10% I-5 hydrogel did not enhance their survival, even though the hydrogel consistently prevented cavity formation and facilitated the creation of ECMs at the lesion epicenter. Our findings indicate that NSC graft survival significantly improved when they were transplanted with a stiffer (16%) I-5 hydrogel compared to a softer (10%) hydrogel. By using an *in vitro* model with precisely controlled mechanical stiffness of the hydrogel substrate, we demonstrated that the mechanical properties of the hydrogel substrate substantially influence NSC behavior and viability accompanied by regulation of cellular elasticity and intracellular Ca^2+^ oscillations in NSCs via an actin polymerization-dependent manner. Additionally, we found that the stiffness-dependent NSC survival in the injured spinal cord was attenuated by Crispr/Cas9-based editing of the *Piezo1* gene in transplanted NSCs, highlighting the critical role of mechanotransduction in NSCs in modulating the viability of grafted NSCs in the injured spinal cord.

Despite the potential of NSC transplantation for repairing damaged spinal cord, poor graft survival remains a significant challenge. Recent studies have shown that using a combination of multiple growth factors and anti-apoptotic molecules at high concentrations could dramatically improve graft survival (*3*). However, it remains to be determined whether this approach could result in undesirable off-target effects, such as allodynia and autonomic dysreflexia, due to nonspecific and unregulated stimulation of host circuits (*1, 58*). Advances in biomaterial engineering have offered potential solutions to enhance post-transplantation cell survival. In this study, while our hydrogel engineered to conjugate the imidazole moiety successfully created eosin-positive matrices at the lesion core, fulfilling its role as a scaffold, NSC survival remained poor when complexed with a 10% hydrogel. Interestingly, when we increased the hydrogel concentration to 16%, which made it nearly five times stiffer, we observed a dramatic improvement in NSC graft survival. In our replication experiment, we confirmed a positive correlation between the stiffness of the I-5 hydrogel stiffness and both the success rate and the volume of surviving NSC grafts. This finding was further supported by an increase in pERK expression in NSCs transplanted with the stiffer hydrogel, indicating enhanced prosurvival intracellular signaling. Importantly, the improved survival was not attributable to differences in hydrogel degradation kinetics, gelation speed, or effects on neuroinflammation, as these parameters remained consistent across different hydrogel concentrations, indicating that the mechanical properties of the hydrogel directly influence NSC survival. Previous studies showed that transplanting NSCs encapsulated in engineered hydrogel scaffolds improves post-graft NSC survival (*59, 60*). Recent studies have also reported that substrate stiffness can regulate the differentiation and the proliferation of NSCs, mainly *in vitro* conditions (*61–63*). However, to the authors’ knowledge, there are no studies directly comparing the effects of different stiffness levels of biomaterial scaffolds on NSC viability, particularly in conditions where NSC survival is compromised, such as in an injured spinal cord. We speculate that the ability of the I-5 hydrogel to create ECM at the lesion epicenter introduces a singular opportunity to examine the potential influence of the mechanical properties of hydrogel on NSC graft survival. Our previous study demonstrated that the hydrogel-created ECM begins to accumulate as early as one week after injection and continues to mature over the course of 4 weeks post-injection (*18*). In addition, it has been reported that CNS tissue, including the spinal cord, becomes softer soon after injury (*36, 64*). Therefore, it is conceivable that the lesioned spinal cord is significantly softer in the days immediately following hydrogel injection than the uninjured spinal cord. During these early days of injection, the viability of NSCs may be highly influenced by the mechanical environment afforded by I-5 hydrogel, which serves as a scaffolding platform before it begins to degrade completely.

Our *in vitro* studies using PAA hydrogel substrates with defined elastic moduli provided mechanistic insights into how substrate stiffness affects NSC behavior. NSCs cultured on stiffer substrates (25 kPa) showed improved adhesion, displayed more complex morphology with extended processes, and exhibited higher viability compared to those on softer substrates (0.2 ∼ 0.5 kPa). Cell-matrix interaction in engineered polymer scaffolds exerts influence on cellular morphology (*65, 66*). Increasing the stiffness of flat hydrogel surfaces led to enhanced spreading of mesenchymal stem cells (*67*), resulting in larger spreading areas, similar to the responses observed with NSCs grown on stiffer substrates in our study. Cellular contact with a stiffer polymer substrate may enhance the activity of the focal adhesion kinase (*19*), which in turn regulates the extent of cellular adhesion to the substrate. In addition, mechanotransduction involves various adaptive cellular responses, including mechanisms of proliferation and cell death (*19*). Our experiment demonstrated that stiffness-dependent viability persists under cellular stress, indicating that a stiffer hydrogel could effectively protect NSCs in the degeneration-prone environment of the injured spinal cord. We measured elasticity and intracellular Ca^2+^ oscillations as markers of active mechanotransduction on stiffer substrates. Both markers were highly dependent on actin polymerization, implying that the mechanotransduction primarily affects the cytoskeletal machinery beneath the cellular membrane. Intracellular Ca2+ oscillations within cells are crucial for various cellular processes, including survival and death (*68*). Consequently, stiffness-dependent Ca^2+^ oscillatory activity may suggest a significant role of mechanotransduction in maintaining cellular homeostasis.

Among various mechanosensitive ion channels expressed in NSCs, Piezo1 emerged as the primary mediator of stiffness-dependent cellular responses in the current study. Inhibition of Piezo1 with GsMTx4 or siRNA knockdown significantly altered NSC morphology and reduced adhesion, specifically on stiffer substrates, mimicking the behavior of NSCs on soft substrates. Electrophysiological recordings confirmed functional Piezo1 channels in NSCs, with pressure-induced inward currents that were blocked by GsMTx4. Importantly, CRISPR/Cas9-mediated Piezo1 gene editing significantly reduced NSC graft survival *in vivo* when transplanted with 16% hydrogel, with success rates dropping from 71.4% to 37.5%. These findings establish Piezo1 as a critical mechanotransducer that senses the stiffness of the surrounding hydrogel environment and regulates NSC survival. Previous studies highlighted the role of Piezo1 in various stem cell functions, including fate differentiation and fate determination (*57, 69*). Activation of Piezo1 in certain tumors helps them evade apoptosis, contributing to the survival and progression of cancerous cells (*70, 71*). Our study appears to be the first report on the role of Piezo1 in regulating the survival of NSCs. Based on a variety of research focused on Piezo1 functions, we can infer that the activation of Piezo1 in transplanted NSCs, caused by the surrounding stiffer hydrogel, can initiate intracellular prosurvival signals, which in turn prevent the loss of NSCs during the early post-transplantation period.

Our findings may have profound implications for cell-based therapies across all CNS injuries, not just spinal cord trauma. Optimizing the mechanical stiffness of hydrogels that encapsulate NSCs or other therapeutic cells could dramatically enhance graft survival, addressing one of the critical challenges in the field of cell-based therapy. Our study also advances our understanding of cellular mechanotransduction in neural repair employing cell transplantation. Regardless of whether hydrogel is used, a suboptimal mechanical environment in the injured CNS may critically undermine the survival of grafted therapeutic cells. Injured spinal cord and brain tissues become significantly softer (*64, 72*), likely due to ECM degradation, increased water content, and loss of myelination, among other factors. This stiffness mismatch between injured and intact tissue likely deprives transplanted cells of essential mechanotransduction-dependent prosurvival signals. The identification of Piezo1 as a key mechanotransducer opens new avenues for enhancing graft survival in these conditions. Pharmacological activation of Piezo1 or genetic engineering to overexpress this channel could potentially boost cell survival in mechanically suboptimal environments. Furthermore, developing hydrogel biomaterials with optimized mechanical properties that minimize stiffness mismatches could maximize Piezo1 activation and subsequent graft survival. Future research should explore the complex interplay between mechanical and biochemical cues in the injury microenvironment to develop comprehensive strategies that enhance both graft survival and functional integration.

## Materials and methods

### Primary neural stem/progenitor cells (NSCs) culture

NSCs were isolated from time-pregnant transgenic SD rats ubiquitously expressing enhanced green fluorescent protein (GFP) as previously described (*6*). Briefly, the spinal cords from E14 (embryo) fetuses of pregnant rats were dissected in a cold Hanks’s Balanced Saline Solution (HBSS, Thermo) containing 1% antibiotics (Thermo) and antifungal (Thermo), and the meninges were carefully removed under the dissection microscope. Subsequently, the spinal cords were mechanically dissociated in a mixture of Accumax (Merck, #SCR006) and DNAse Ⅰ (Sigma, #10104159001, 20 μg/ml) and the dissociated cells were passed through a 40 μm cell strainer to remove debris followed by centrifugation for 5 min at 1000 RPM. The cells were resuspended in StemPro**^®^**NSC SFM culture media (Thermo, #A1050901) containing bFGF (20 ng/ml) and EGF (20 ng/ml), and plated into a non-coated Petri dish (SPL, #10093) to grow as neurospheres in a humidified atmosphere with 5% CO_2_. The culture medium was changed every other day.

### Preparation of NSC-hydrogel complex

In our previous studies (*18, 73*), we have provided a comprehensive outline of the synthesis procedure for the I-5 hydrogel. Briefly, poly(dichlorophosphazene) (10 g, 86.29 mmol) was sequentially reacted with isoleucine ethyl ester (23.81 g, 121.67 mmol), 2-aminoethanol (1.56 g, 25.5 mmol), and aminopolyethylene glycol (56.29 g, 75.06 mmol) to poly(organophophazene). Subsequently, the hydroxyl group of 2-aminoethanol was esterified using succinic anhydride (2.76 g, 27.58 mmol) to create a hydrolysable ester linkage and a terminal carboxylic acid group. Finally, I-5 was synthesized by conjugating 1-3 aminopropylimidazole (1.45 g, 11.54 mmol) to the carboxylic acid group. To prepare the NSC-hydrogel complex, neurospheres were collected into 15 ml conical tubes by centrifugation for 1 min at 1000 RPM on the day of transplantation. Cell pellets were incubated in Accumax**^®^** for 3 min at RT, followed by gentle mechanical dissociation into single cells. After centrifugation, cell pellets were resuspended at 5 × 10^5^ cells/μl in a culture medium and kept on ice until complexation with I-5 hydrogel. Dissociated NSCs in a culture medium were mixed with an appropriate volume of I-5 hydrogel in a solution state (20% stock polymer solution) to achieve final polymer concentrations of 10%, 13%, and 16%. To determine whether adding an exogenous growth factor or an extracellular matrix protein could improve graft survival, we included insulin-like growth factor-1 (R&D systems, #4326-RG, 2 μg/ml) or laminin (Thermo, #23017015, 3 μg/ml) to the culture medium prior to the complexation with I-5 hydrogel.

### Animals and surgical procedures

All animal protocols were approved by the Institutional Animal Care and Use Committee (IACUC) of Ajou University School of Medicine, South Korea. Adult female Sprague– Dawley (SD) rats (8 weeks) weighing 200–230 g (Orient Bio, South Korea) were used for all animal experiments. The SCI model was created by using the Infinite Horizon Impactor (Precision Systems and Instrumentation), as described in our previous studies (*18, 43*). Briefly, the animals were deeply anesthetized by intraperitoneal injection of ketamine (80 mg/kg) and xylazine (10 mg/kg) mixtures. The hindlimb pedal withdrawal reflexes were examined to determine adequate anesthesia level before starting surgery. A longitudinal dorsal incision was created between T7-T11 spinal vertebrae, followed by a laminectomy at the level of T9. Then, the subjects underwent a contusive SCI with a force of 200 kdynes, which replicates the clinically relevant pathophysiological events following SCI. Following surgery, animals were placed in a warming incubator at 37°C until they were fully awake. Postoperative care included expressing the bladder twice daily until natural urination resumed. NSC transplantation was performed one week after SCI. The total number of transplanted NSCs per animal was 1 × 10^6^ cells in 10 μl of NSC-hydrogel complex prepared as described above. The NSC-hydrogel complex was loaded into a 26G Hamilton syringe (Hamilton #80030) and kept on ice for at least 1 h until transplantation. It was crucial to maintain a temperature of 4℃ to ensure the solution state of the complex before transplantation. After re-exposing the lesioned spinal cord, 10 μl of the NSC-hydrogel complex was removed from the ice and injected into the lesion epicenter over 1 min without any delay. The syringe was then kept in place for 1 min to prevent leakage. Postoperative care was provided in the same way after the transplantation procedure.

### Assessment of physical properties of I-5 hydrogel

Rheological measurement of the 10% and 16% I-5 hydrogel was conducted using a rheometer (MSC 102, Anton Paar) over a temperature range of 5 to 60 °C at an oscillation frequency of 1Hz and a strain of 10 %. Storage and loss moduli were obtained using the instrument software. Young’s modulus (E) was estimated using the measured storage modulus (G′) and loss modulus (G″) from rheology data. The complex shear modulus (G*) was first calculated using the equation: 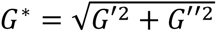. Assuming isotropic material behavior, Young’s modulus was then derived from the shear modulus using the standard relationship: E = 2G(1 + ν) where G≈G’ (complex modulus from rheometer), ν = poisson’s ratio (for soft materials like hydrogels, assume near 0.5). To assess the gelation speed, hydrogel was prepared by adding 0.2 mg/ml of Nile Red (Sigma) to achieve final concentrations of 10 % and 16 %. The mixture underwent 1h of sonication and was loaded into a 31G needle syringe. Meanwhile, 20 mL of water in a vial was heated to 37°C, and the hydrogel was subsequently injected into it. The gelation speed was visually monitored and compared between the two different concentrations. For the in vivo hydrogel dissolution test, all animal experiments were approved by the IACUC at the Korea Institute of Science and Technology (KIST). Six-week-old female Balb/c mice (Orient Bio, South Korea) were anesthetized using 3% isoflurane in a mixture of oxygen and nitrogen. The hydrogel was mixed with 0.2 mg/ml nile red to achieve concentrations of 10 % and 16 % and sonicated for 1 h. Then, 50 µl of hydrogel samples were loaded into a 31G needle syringe and injected subcutaneously into the dorsal back skin of mice, with the left side receiving the 10% and the right side receiving the 16%. The retention of hydrogel was monitored at prearranged time points using an IVIS spectrum imaging system (Caliper, USA).

### Cryogenic scanning electron microscopy (cryo-SEM)

The cryo-SEM experiments were conducted using a Quanta 3D FEG microscope (FEI, Netherlands) with an Alto 2500 cryo-transfer system (Gatan, UK). The I-5 hydrogel was complexed with 1× 10^6^ cultured NSCs at a concentration of 10 wt%. The samples were rapidly frozen in liquid nitrogen and moved to a freezing chamber pre-evacuated to 10^-5^ mbar at −190°C. Metal deposition was achieved via sputtering with a 3-mA current, and the samples were then transferred to a microscope chamber also pre-evacuated to 10^-5^ mbar at - 190°C. Cryo-SEM images were taken using a 5-keV beam at 11.8 pA.

### Tissue processing and histological experiments

Animals were deeply anesthetized using a mixture of ketamine (80 mg/kg) and xylazine (10 mg/kg), followed by intracardiac perfusion with cold PBS and 4% paraformaldehyde (PFA) in 0.1 M phosphate buffer. The tissue covering the lesion epicenter of the spinal cord (± 0.9 mm rostral and caudal) was carefully dissected out, and then post-fixed with 4% paraformaldehyde for 6 h at 4°C, followed by cryoprotection in a series of 10% and 30% sucrose solutions for 5 days. The spinal cord tissues were embedded in an optimal cutting temperature compound and then longitudinally sectioned at a thickness of 20 μm using a #CM1900 cryostat (Leica). To assess lesion cavities, tissue sections were immersed in a staining solution composed of 240 ml of 0.2% Eriochrome Cyanine RC (ECRC) and 10 ml of 10% FeCl3·6H2O in 3% HCl for 5 min, and then washed with running tap water for 5 min. Next, the tissues were treated with 1% NH_4_OH for differentiation and counterstained with eosin solution to make lesion cavities induced by hydrogel injection more discernible. Picrosirius red staining (abcam, ab150681), which visualizes total collagen fibrils by binding to the aligned basic amino acids on the collagen molecule regardless of the type [32], was performed to evaluate the fibrotic environment within the hydrogel-created ECM according to the manufacturer’s instruction. For immunohistochemical staining, tissue sections were blocked in a 10% normal goat serum and 0.3% Triton-X in PBS for 1 h at RT. Then, the primary antibodies were incubated in a blocking solution at 4 °C overnight. The following primary antibodies were used: chicken anti-GFP (Abcam, ab13970, 1:3000), rabbit anti-fibronectin (Sigma, F3648, 1:300), mouse anti-fibronectin (Millipore, MAB1940, 1:100), rabbit anti-GFAP (DAKO, Z0334, 1:2000), chicken anti-GFAP (Abcam, ab4674, 1:5000), rabbit anti-phospho p44/42 MAPK (ERK1/2, Cell signaling, 4370S, 1:200), rabbit anti-Iba1 (Wako, 019-19741, 1:1000), rabbit anti-5-hydroxytryptamine (5-HT, Immunostar, 20080, 1:5000), mouse anti-synaptotagmin (Millipore, MAB5200, 1:500), mouse anti-nitrotyrosine (Abcam, clone HM.11, 1:300), and rabbit anti-Tubb3 (Sigma, T2200, 1:2000). After washing off unbound primary antibodies with PBS three times for 5 min each, the samples were incubated with the appropriate fluorescent Alexa Fluor secondary antibodies (Thermo) for 2 h at RT. Following the antibody washings, tissue sections were mounted with an anti-fade DAPI solution (Vector). Images were then obtained using an LSM 800 confocal laser-scanning microscope (Carl Zeiss), an Axio Scan.Z1 slide scanner (Carl Zeiss), or an LSM 900 confocal laser-scanning microscope with Airyscan super-resolution (Carl Zeiss).

### Analyzing images of histological samples

To evaluate the graft volume of NSCs *in vivo*, three consecutive spinal cord tissues (300 μm interval between sections) covering the lesion epicenter were chosen for each animal. Images were acquired from the Zeiss Axio Scan.Z1 slide scanner (Carl Zeiss) microscope at a 200× magnification. The lesion epicenter was delineated by using GFAP immunoreactivities. The boundaries along surviving grafts with discernible GFP signals were manually delineated using the Zen Lite 3.0 spline contour tool (Carl Zeiss), and then the total estimated volume was calculated using Cavalieri’s principle as described in our previous study (*6*). Graft success was determined by the presence of distinguishable grafts within the lesion epicenter in at least one tissue section. To evaluate the expression of phosphorylated extracellular signaling-related kinase 1/2 (phospho-Erk1/2) in grafted neural stem cells (NSCs), images were captured using a confocal laser-scanning microscope with a 20× objective lens and consistent exposure settings across all groups. Four regions of interest (ROIs) from three sections per animal were randomly placed within the lesion epicenter to cover the noticeable GFP positive engraftment. The number of NSCs doubly positive for GFP and phospho-Erk1/2 was manually counted, and the percentage of the double positive cells out of the total GFP-positive cells within ROIs was calculated. The percentage of NSCs that differentiated into either neurons or astrocytes was determined using the same method. To assess the expression of fibrotic matrix formation at varying hydrogel concentrations, images of fibronectin and Picrosirius red staining were obtained using Zeiss Axio Scan.Z1 slide scanner (Carl Zeiss) with 200× magnification. Two regions of interest (500 μm × 500 μm) were selected from the brightest areas within the lesion epicenter in each section, and four regions of interest per animal were analyzed to calculate an average value. To quantify the 5-HT axonal ingrowth into the hydrogel-created matrix, a single spinal cord section containing the largest NSC grafts was selected. The length of 5-HT axons within hydrogel-induced matrix was manually measured using the NeuronJ plug-in in the ImageJ software. Co-localization of doubly positive 5-HT axons with NSC grafts was manually counted using the Zen Lite software (Carl Zeiss). To create a 3D reconstruction image visualizing synaptic contacts between host axons and grafted NSCs, we performed triple immunofluorescence staining using antibodies against GFP, 5-HT, and synaptotagmin. Images were captured using a laser-scanning microscope with a 63× objective lens and imported into IMARIS 9.0 software for 3D reconstruction. Background subtraction and the surface rendering method were used to create the 3D images following the instruction manual.

### NSC adhesion and spreading assays on substrates covered with hydrogels of varying stiffness

Cell culture vessels coated with polyacrylamide (PAA) hydrogel, featuring different elastic modulus levels, were purchased from Matrigen (http://matrigene.com). PAA hydrogel was bound to either coverslips in a 12-well polystyrene plate (#SW12-EC, Matrigen) or glass-bottom coverslips in a confocal dish (#SV3520-EC, Matrigen). Hydrogel coverslips were pre-coated with poly-D-lysine (Sigma #P6407, 0.01 mg/ml) for 18.5 h at 37 °C and gently washed with distilled water three times followed by drying for 1 h at RT. Cultured primary NSCs ubiquitously expressing GFP grown as neurospheres were collected into a 15 ml conical tube and mechanically dissociated into single cells followed by plating them onto the hydrogel coverslips in a DMEM + 3% FBS medium for 24 h. Subsequently, cold 4% PFA was used to fix the cells for 15 min at RT, and then washed three times with PBS followed by blocking steps for 1 h with 10% normal goat serum containing 0.3% triton-X in PBS for immunostaining with following antibodies: chicken anti-GFP (abcam, #ab13970; 1:2000), rabbit anti-phospho-ERK1/2 (Cell signaling, 4370S, 1:200). To visualize actin filament, Alexa Fluor Phalloidin (Thermo, #A12381) was used according to the manufacturer’s instruction. For pharmacological inhibition of mechanosensitive ion channels, cultured NSCs were treated with GsMTx4 (Piezo1 inhibitor, abcam, ab141871; IC50 5 µM), Pico145 (TRPC1 inhibitor, Medchem, HY-101507; IC50 1.3 nM), and Amiloride hydrochloride (TRPP2 inhibitor, Medchem, HY-B0285A; 2.6 µM) in a DMEM + 3% FBS medium for 24 h. Dosages for various drug treatments were determined based on the IC50 value of each reagent. For quantitative evaluation of NSC adhesion on hydrogel substrate with different stiffness, three regions of interest (ROIs) per coverslip were randomly placed, and images were obtained using an LSM 800 confocal laser-scanning microscope at 100× magnification with identical exposure settings across all the experimental groups. The number of GFP+/DAPI+ cells was manually counted using Zen lite software, and the counts from all images were averaged to acquire one value per replicate. For the quantitative assessment of morphological changes in NSCs upon spreading onto the hydrogel substrate, we utilized the analysis methods outlined in previous publications (*74, 75*) with a minor modification. Images from five ROIs per coverslip were acquired at 200× magnification. The cell outlines in the images were manually traced using ImageJ. The area and perimeter were then obtained using the measurement function in ImageJ. The total spreading area and perimeter were divided by the number of cells to obtain the average spreading area and perimeter per cell.

### Live/dead cell survival assay in vitro

Neurospheres were collected and dissociated into single NSCs and the cells were plated into Poly-D-Lysine pre-coated hydrogel coverslips in a DMEM+3% FBS medium. NSCs were treated with or without sodium arsenite (NaAsO_2_, Merck, #106277, 5 μM) to induce oxidative stress, mimicking the microenvironment in the injured spinal cord. After 24 h, cells were washed with HBSS and then incubated with the LIVE/DEAD™ Viability/Cytotoxicity Kit (Invitrogen, #L3224) according to the manufacturer’s instructions. Briefly, cells were treated with 0.5 μM of calcein AM (495/517 nm) and 1 μM of ethidium homodimer-1 (528/617 nm) for 20 min. Live cells convert nonfluorescent calcein AM into a green fluorescent calcein by hydrolyzing acetoxymethyl ester. Ehidium homodimer-1 enters cells with damaged membranes, thereby producing a red fluorescence. Subsequently, images were captured from five randomly selected ROIs on a coverslip using a fluorescence microscope with 100× magnification. The images were then converted to 8-bit and adjusted using identical thresholds with post-processing using the Watershed function. The number of dead cells (red) was automatically quantified using the ImageJ software with an automatic cell counting module. To evaluate the potential damage to NSCs during an injection procedure, cell suspensions were loaded into a 26G Hamilton syringe (Hamilton #80330) and then injected into a 24-well culture plate using the same parameters used for *in vivo* injection (10 μl/min, 1 min rest). After 30 min, the live/dead assay was conducted as described above using cells obtained before injection and those injected using the 26G Hamilton syringe.

### Calcium imaging

For spontaneous intracellular Ca^2+^ imaging, glass bottom coverslips or hydrogel-bound coverslips in a confocal dish (Matrigen, CA, #SV3520-EC) were pre-coated with Poly-D-Lysine for 18 h in a 37 °C incubator. Dissociated NSCs were plated and incubated for 24 h, and then the culture medium was washed out with HBSS. Subsequently, NSCs were loaded with 2 μM of membrane-permeable calcium dye Fluo-4 AM (Invitrogen, #F14201) in Hanks balanced salt solution (HBSS, Invitrogen) for 30 min in a 37 °C incubator, according to the manufacturer’s instruction. The cells were washed three times with HBSS and then underwent complete de-esterification of the dye for an additional 10 min at RT. Within 30 min of calcium dye loading, calcium imaging was performed in a bath solution containing 0.63 mM of calcium chloride, 0.245 mM of magnesium chloride, 0.2 mM of magnesium sulfate, 2.65 mM of potassium chloride, and 68.5 mM of sodium chloride using a LSM800 laser scanning microscope. Spontaneous Ca^2+^ oscillations were monitored using time-lapsed images at 1-sec intervals for a duration of 1 min. For quantitative evaluation of Ca^2+^ oscillations, images were captured at 400× magnification. Five regions of interest (ROIs) were randomly selected, and 10 cells per image were analyzed, totaling 50 cells per group, using ImageJ with the Time Series Analyzer plugin. Three independent experiments were performed. Fluorescent signals were normalized with the baseline and presented as a ratio of the mean change in fluorescence (F_max_/F_0_). The baseline fluorescent signal (F_0_) was obtained from the first or last 10 sec when the fluorescent level was the lowest. Calcium-responsive cells were defined as cells showing more than a 20% increase over the baseline level, while cells showing less than 20% of the baseline were considered non-responsive. For the drug pre-treatment experiment, cells were treated with 20 μM of cytochalasin D (Sigma, #C8273) for 30 min prior to loading of Fluo-4 AM. For the experiment with Yoda1 (Tocris, 5586, 100 μM), baseline Ca^2+^ activity was recorded for the first 30 sec and applied the Yoda1 during the imaging.

### Electrophysiology

Whole-cell patch-clamp recordings were conducted at RT. The recording chamber was continuously perfused with a HEPES extracellular recording solution containing 135 mM NaCl, 5.4 mM KCl, 1.8 mM CaCl_2_, and 5 mM HEPES. The pH was adjusted to 7.4 with KOH and HCl, and the osmolality was set to 308 ± 2 mmol/kg. Voltage-clamp recordings utilized Cs^+^ solutions, with the Cs^+^ intracellular solution containing 90 mM Cs Methanesulfonate, 48.5 mM CsCl, 2 mM MgCl_2_, 5 mM Cs-EGTA, 2 mM NaATP, and 5 mM HEPES. The pH of the Cs^+^ solution was adjusted to 7.2 with KOH, and the osmolality was set to 307 ± 2 mmol/kg. Patch pipettes were pulled from borosilicate glass using a vertical micropipette puller (PC-100, World Precision Instruments) and flame polished to achieve resistances of 5-7 MΩ. Signals were recorded using a Multiclamp 700B amplifier (Molecular Devices, Union City, CA), filtered at 2 kHz, and sampled at 10 kHz. A Picospritzer III delivered air pulses at 7 psi pressure to apply mechanical stimulation to the cells. The solution used in the pipette for mechanical stimulation had the same composition as the HEPES extracellular recording bath solution. Data were collected at a sampling frequency of 10 kHz and analyzed offline using Clampfit software. All group data are presented as mean ± SEM, and statistical comparisons were performed using a two-way ANOVA. All drugs used in the electrophysiology experiments were purchased from Sigma-Aldrich.

### Measurement of elastic modulus by atomic force microscopy

Cellular elasticity was measured using atomic force microscopy (AFM) (Nano N8 Neos; Bruker Corporation, Germany) with force-distance (FD) curve measurement in liquid condition (*76*). The FD curve was measured by using an AFM probe (Cont GD, Budget Sensors Inc, Bulgaria) composed of an Au-coated cantilever and Si-based conical tip. Detailed dimensions of the probe were as follows: resonance frequency, 13 kHz (± 4 kHz); force constant, 0.2 N/m (0.07-0.4 N/m); cantilever length, 450 µm (± 10 µm); cantilever width, 50 µm (± 5 µm); cantilever thickness, 2 µm (± 1 µm); tip height, 17 µm (± 2 µm); tip radius < 10 nm. The half-cone angle of the conical tip along the cantilever axis ranged from 20 ∼ 22.5°. The load force was set at ≤ 10 nN to minimize damage to the cell membrane, and the loading rate of the probe was approximately 1 µm/s. FD curve measurement was performed at 10 points per cell for 20 to 31 cells of each condition. The cellular elasticity was measured around the nucleus, where the cell thickness was approximately 1 μm or more, to avoid stiffness effects due to the nucleus and matrix. NSCs were seeded on the hydrogel-bound coverslips and incubated for 24 h in DMEM + 3% FBS and then the cells fixed with 3.7% paraformaldehyde for 15 min at RT followed by washing three times with PBS. The samples were submerged in PBS at 4 °C until AFM measurement.

### RNA extraction and qPCR

Total RNA was isolated from dissected tissues or cultured cells using Trizol (Invitrogen) according to the manufacturer’s instructions. RNA quantity and purify were analyzed using a NanoDrop Life Spectrophotometer (Thermo Fisher Scientific) at 260 nm. One microgram of RNA was reverse transcribed to cDNA using the manufacturer’s protocol (Cellsafe, #CDS-100). One microliter of cDNA was added to SYBR green Master Mix (Takara) containing 10 pM of the primer pairs listed in the Table S3. Quantitative real-time PCR was performed using a 7500 Real-Time PCR system (Applied Biosystems, Foster City, CA, USA) according to the manufacturer’s protocol. The cycling conditions were 95 °C for 30 sec and 54.8∼61.5 °C for 5 sec for a total of 40 cycles. Melting curves were generated after the final extension step, and Ct values were calculated using the Applied Biosystems 7500 software. The Ct values of the target genes were normalized to the 18S rRNA housekeeping gene as an internal control.

### Gene silencing experiment with siRNA

Piezo loss of function experiments were performed using small interfering RNA (siRNA, Dharmacon, #361430). Briefly, five million dissociated NSCs were transferred into a nucleofector cuvette containing Nucleofector® solution (Lonza, Rat Neural Stem Cell Nucleofector™ Kit, #VPG-1005) with 2 μM of siRNA Piezo1 or scrambled siRNA. Subsequently, the cuvettes were inserted into a Nucleofector® 2b (Lonza) using Rat Neural Stem Cell Nucleofector® Kit (Cat No: VPG-1005) followed by electroporation. The cells were then allowed to recover for 24 h in a culture plate before the subsequent experiments. ***CRISPR/Cas9-mediated Piezo1 editing in NSCs***

The CDS region of rat *Piezo1* gene was screened for sgRNA using the Cas-Designer tool. (http://www.rgenome.net/) (*77*). Small guide RNAs (sgRNAs) were selected based on filtering with mismatch criteria of 1,0,0. These sgRNAs were synthesized *in vitro* and transfected into C6 gloma cells (ATCC, CCL-107) for screening. After transfection, lead sgRNA was selected based on their high gene editing activity, as determined by targeted deep sequencing, and by the downregulation of target gene expression at the mRNA level. For targeted gene editing in NSCs, sgRNA and Cas9 protein were introduced via electroporation using a 4D-Nucleofector (Lonza, P3 Primary Cell 4DNucleofector X Kit) as a ribonucleoprotein (RNP) complex. Briefly, the RNP complex was formed by mixing 1 μg Cas9 protein with 4 μg sgRNA and incubating at room temperature for 10 min. This complex was then electroporated into 4 × 10^5^ rat NSCs using 20 μl of primary P3 buffer (Lonza). At least 72 h after transfection, genomic DNAs (gDNAs) were extracted using a genomic DNA extraction Kit (Favorgen) according to the manufacturer’s protocol. To identify gene editing efficiency and the exact sequence edited, targeted deep sequencing was performed using gDNAs as previously described (*78*). Briefly, gDNAs were amplified using specific primer sets. Paired-end sequencing was then performed using an Illumina MiSeq (Primer used in this study are described in Table S3). Indel frequency and sequence edits were analyzed using the online Cas-Analyzer tool (www.rgenome.net).

### Statistical analysis

Statistical analyses were performed using GraphPad Prism software (version 8.0.2). Unpaired Student’s t-test (two-tailed) was used to compare the mean values of two groups. One-way ANOVA followed by Tukey’s *post hoc* analysis was used for the mean values of three or more groups. Two-way ANOVA followed by Bonferroni’s *post hoc* analysis was performed in experiments with two independent variables.

## Funding

Korea Institute of Science and Technology 2E33781 (YMK)

Pan-ministry full-cycle medical device research and development program RS-2023-00244748 (YMK, BGK)

National Research Foundation of Korea RS-2021-NR061536 (YMK)

National Research Foundation of Korea 2021M3E5D9021367 (SCS)

National Research Foundation of Korea RS-2019-NR040055 (BGK)

National Research Foundation of Korea RS-2021-NR056919 (BGK)

## Author contributions

Conceptualization: HHP, DHH, YMK, SCS, BGK

Methodology: HHP, KSK, JYL, JMP, YMK, BGK

Investigation: HHP, YK, BSJ, SG, DHH, YS, SAJ, HGS, HJK, SL, SK, KIL

Visualization: HHP, YK, BSJ, SAJ, HSK

Supervision: KSK, JYL, JMP, YMK, SCS, BGK

Writing-original draft: HHP, YK, AE, HJK, BGK

Writing—review & editing: JYL, JMP, YMK, BGK

## Data and materials availability

All data from the authors are available upon reasonable request.

