## Supplementary material for "Mechanical environment afforded by engineered polymer hydrogel critically regulates survival of neural stem cells transplanted in the injured spinal cord via Piezo1-mediated mechanotransduction": supple figure and tables

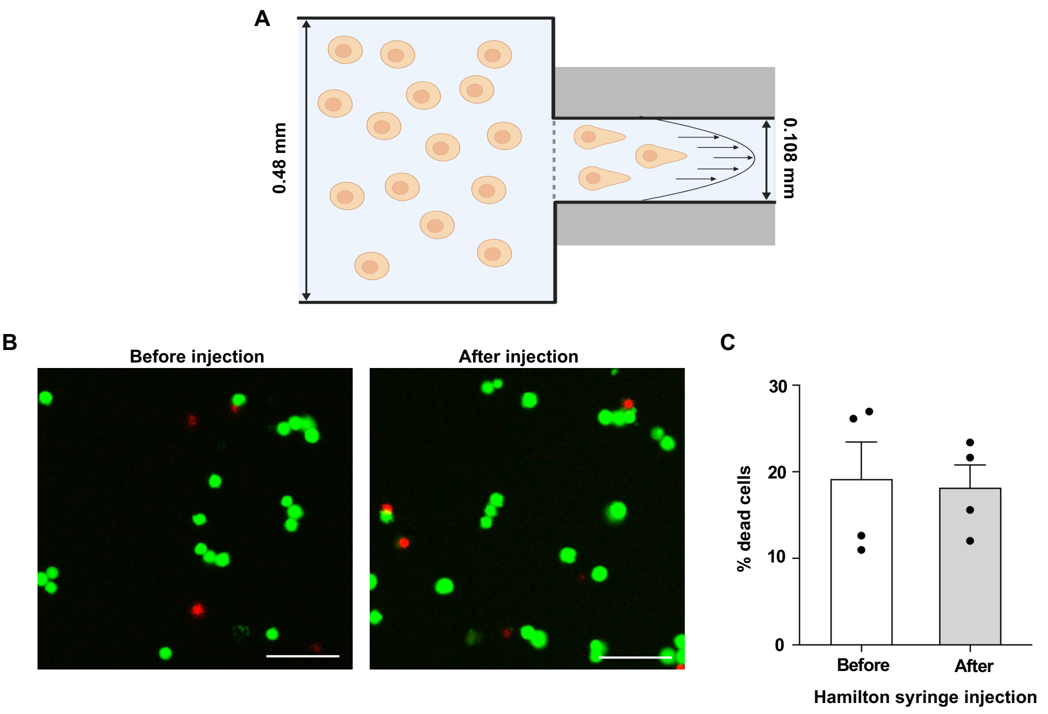


**Figure S1. Syringe needle pressure does not influence the NSC viability**

**(A)** Schematic diagram of the cell viability test caused by syringe needle pressure during injection. **(B)** Representative images of cell survival assay before and after injection. Neurospheres were enzymatically dissociated into single cells, loaded into a Hamilton syringe, and injected into culture medium, mimicking the *in vivo* transplantation study. Live and dead cells were stained using LIVE/DEAD™ Viability/Cytotoxicity Kit. Live cells labeled with Calcein-AM (green) and dead cells labeled with Ethidium Homodimer-1 (red). Scale bar = 100 μm. **(C)** Quantitative graph showing the percentage of dead cells before and after needle injection. Each dot represents one experiment, with averaging of three injection attempts per experiment. Four independent experiments were conducted. Error bars represent SEM.

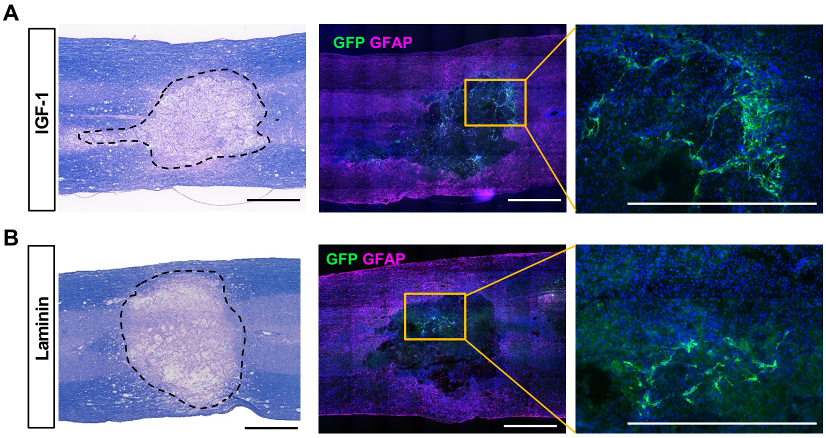


**Figure S2. Combination of a growth factor or ECM protein does not improve survival of NSCs transplanted with hydrogel**

**(A)** Representative images of longitudinal spinal cord sections obtained from animals transplanted with NSCs in 10% hydrogel combined with insulin growth factor-1 (IGF-1) or laminin. The samples were collected at 4 weeks post-transplantation. Spinal cord sections were visualized using eriochrome cyanine and eosin staining (left panel). GFP indicates surviving NSC grafts (green) and GFAP demarcates the lesion areas (magenta, middle panel). Boxed regions are magnified on the right side. Scale bar = 1000 μm.

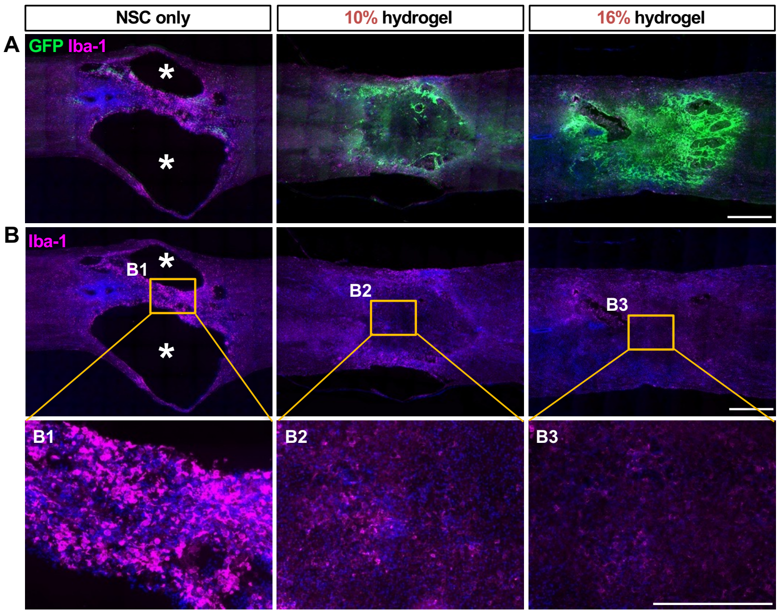


**Figure S3. No difference in the extent of inflammatory reactions between 10% and 16% hydrogel groups**

**(A-B)** Representative images of longitudinal spinal cord sections obtained from animals that received only NSCs or those complexed with 10% or 16% hydrogel. The samples were collected at 4 weeks post-transplantation. Neuroinflammation was evaluated by Iba-1 immunostaining. Boxed regions correspond to magnified views below (B1-B3). Asterisks indicate cystic cavity spaces. Scale bars = 500 μm.

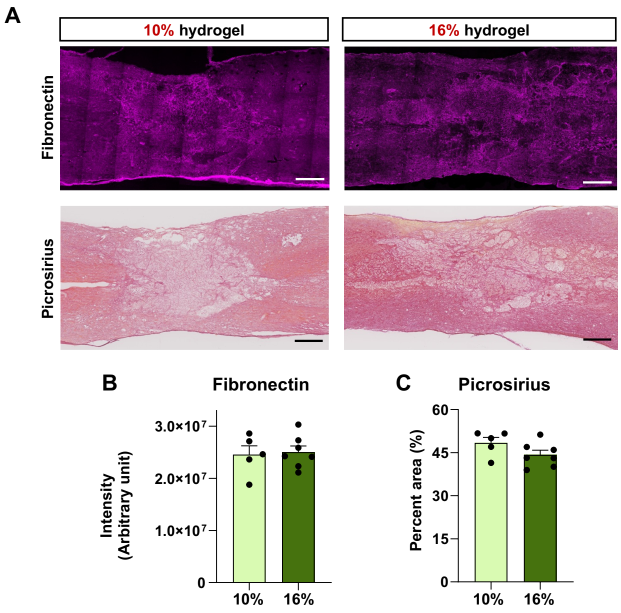


**Figure S4. A difference in hydrogel concentration does not influence the extent of fibrotic matrix formation**

**(A)** Representative images of longitudinal spinal cord sections showing fibronectin staining (top panel) and picrosirius staining for total collagen matrix (bottom panel). No significant differences in fibrotic extracellular matrix formation were observed between groups. **(B)** Quantitative graphs comparing two groups. N=5 and 7 for 10% and 16% hydrogel groups. Error bars represent SEM. Scale bars = 500 μm.

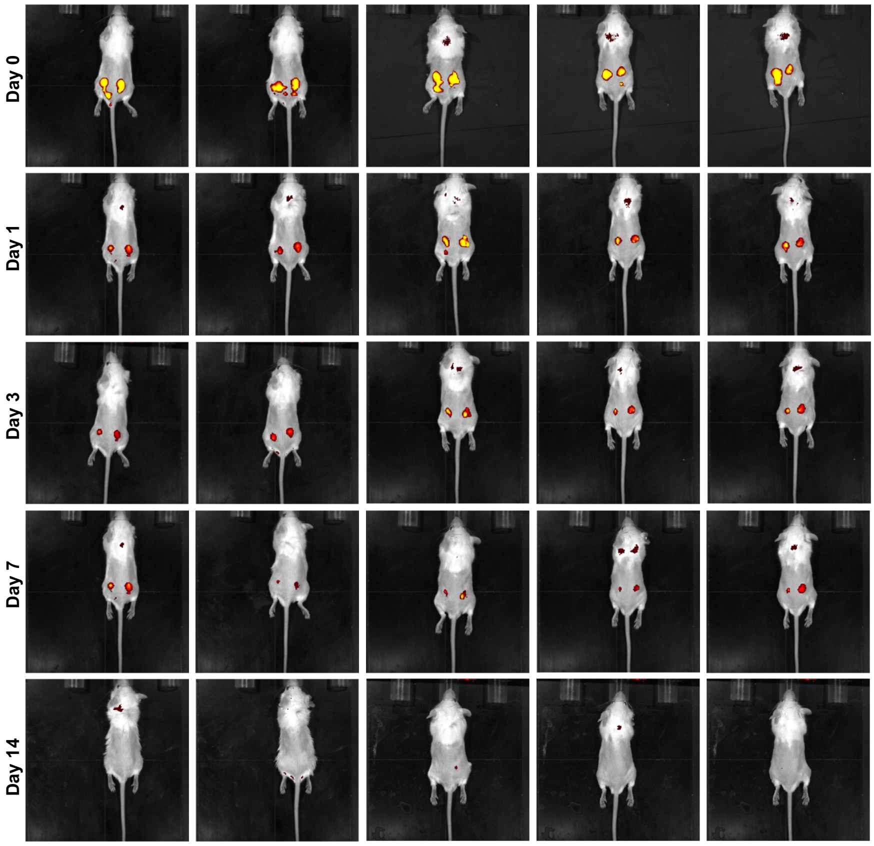


**Figure S5. Hydrogel degradation kinetics with different concentrations of hydrogel**

Representative IVIS spectrum imaging system (Caliper, USA) images showing the mass loss of 50 μl of Nile Red-labeled hydrogel over 0, 1, 3, 7, and 14 days following hydrogel injection. A total of 50 μl of hydrogel samples were loaded into a 31G needle syringe and injected subcutaneously into the dorsal back skin of six-week-old Balb/c mice, with the left side receiving the 10% and the right side receiving the 16%. The left and right panels correspond to 10% and 16% hydrogels in the same animal subject. A total of five animals were monitored during the experimental period.

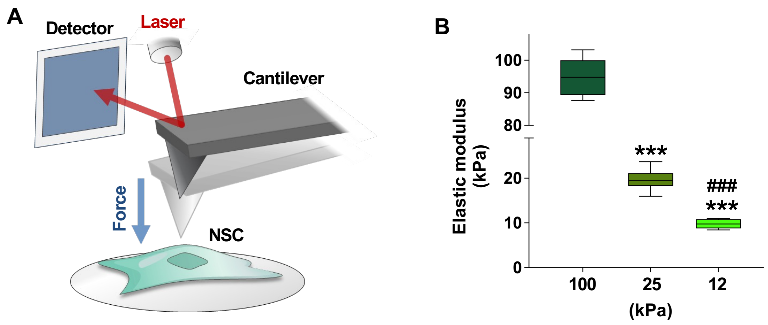


**Figure S6. Measurement of the hydrogel substrate stiffness using atomic force microscopy (AFM)**

**(A)** A schematic diagram illustrating the experimental setup of the AFM to measure the elastic modulus of the NSC plasma membrane. **(B)** Quantitative graph measuring the elastic modulus of the hydrogel culture substrates with varying degrees of mechanical stiffness. N = 12, 25, and 5 cells for 100 kPa, 25 kPa, and 12 kPa groups, respectively. *** indicate $p < 0.001$ compared to the 100 kPa group, and ### indicate $p < 0.001$ compared to the 25 kPa group by one-way ANOVA followed by Tukey's *post hoc* analysis. Error bars represent SEM.

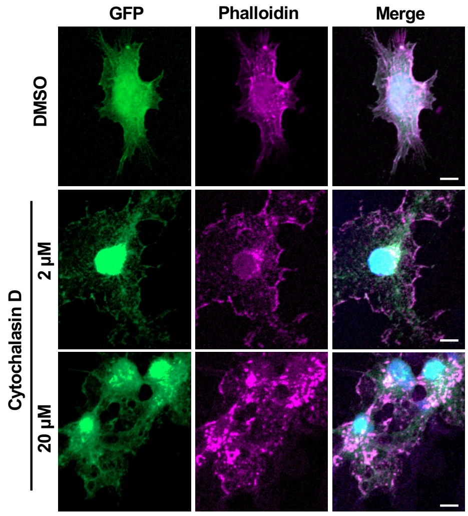


**Figure S7. Cytochalasin D treatment destabilizes filamentous actin cytoskeleton in NSCs in a dose-dependent manner**

Representative images of GFP-positive NSCs cultured on 25 kPa hydrogel substrate and treated with 2 or 20 μM cytochalasin D (actin destabilizer) for 30 min before fixation. Actin filaments (F-actin) were visualized using Alexa Fluor 594-Phallodin. Scale bars = 5 μm.

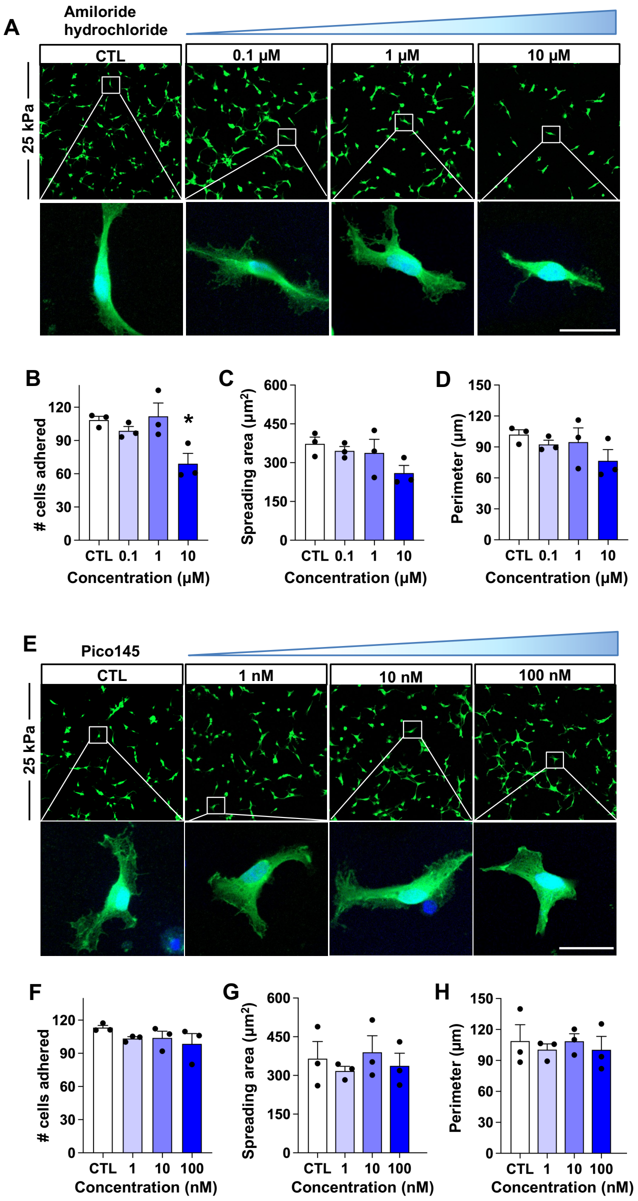


**Figure S8. Neither TRPC1 nor TRPP2 channels influence NSC adhesion on a rigid hydrogel substrate.**

**(A)** Representative images of NSCs cultured on the 25 kPa hydrogel substrate treated with TRPP2 inhibitor (amiloride hydrochloride) for 24 h. **(B-D)** Quantitative graphs comparing the number of cells adhered **(B)**, areas of cell spreading **(C)**, and the perimeter of cell boundary **(D)**. Each dot represents an independent culture replicate, with each replicate being a measurement from one coverslip. **(E)** Representative images of NSCs cultured on 25 kPa hydrogel substrate treated with TRPC1 inhibitor (Pico145) for 24 hrs. **(F-H)** Quantitative graphs comparing the number of cells adhered **(F)**, areas of cell spreading **(G)**, and the perimeter of cell boundary **(H)**. Each dot represents an independent culture replicate, with each replicate being a measurement from one coverslip. Error bars represent SEM. Scale bars = 20 μm.

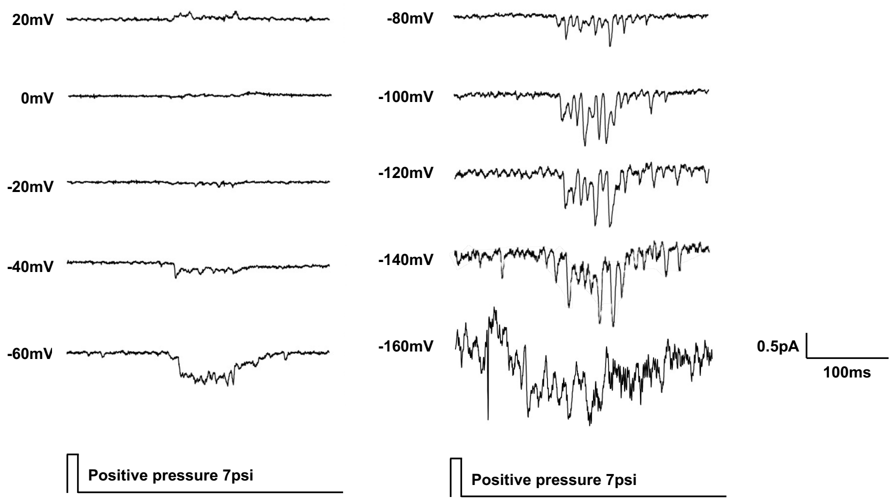


**Figure S9. Representative raw current traces from NSCs in response to pressure stimulation**

Raw current traces were obtained under a voltage clamp mode at varying holding voltages. Short air pulses were applied to NSCs lasting 10 ms with 7 psi pressure using a Picospritzer device.

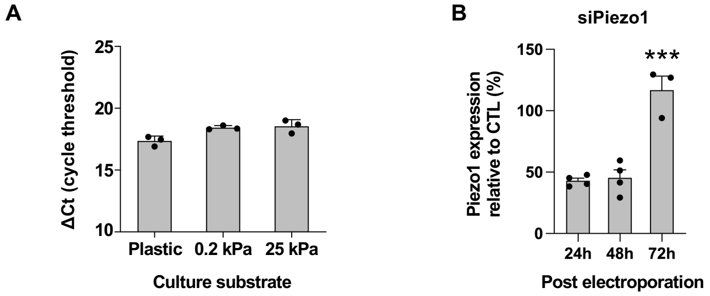


**Figure S10. Validation of Piezo1 knockdown in NSCs**

**(A)** Quantitative PCR (qPCR) analysis of *Piezo1* mRNA expression in NSCs cultured on substrates of varying stiffness. N = 3 independent cultures. **(B)** Validation of Piezo1 knockdown in NSCs by electroporation. *** indicates $p < 0.001$ by one-way ANOVA followed by Tukey's *post hoc* analysis. N = 3 independent cultures. Error bars represent SEM.

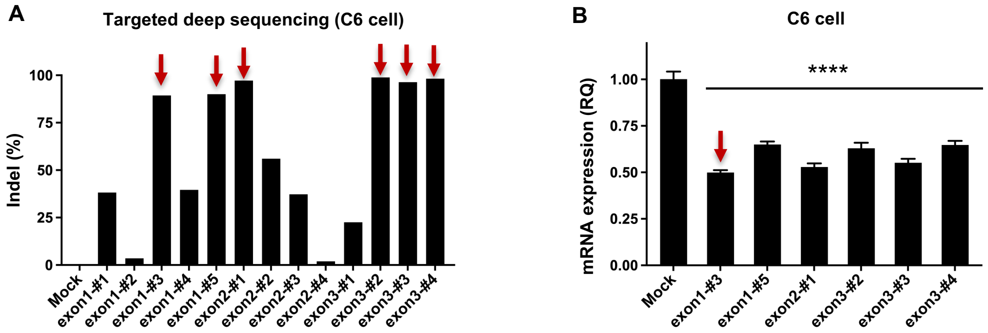


**Figure S11. Screening of CRISPR/Cas9 associated gRNA sequences targeting rat Piezo1**

**(A)** Targeted deep sequencing-based gene editing efficiency of 13 candidate sgRNA sequences in C6 glioma cells. Arrows indicate sequences with Indel frequency higher than 90%. **(B)** Quantitative RT-PCR measuring the levels of Piezo1 in C6 glioma cells that were edited using 6 sgRNA sequences that showed the highest Indel efficiency. An arrow indicates the sequence showing the largest reduction of Piezo1 mRNA level. Error bars represent SEM.

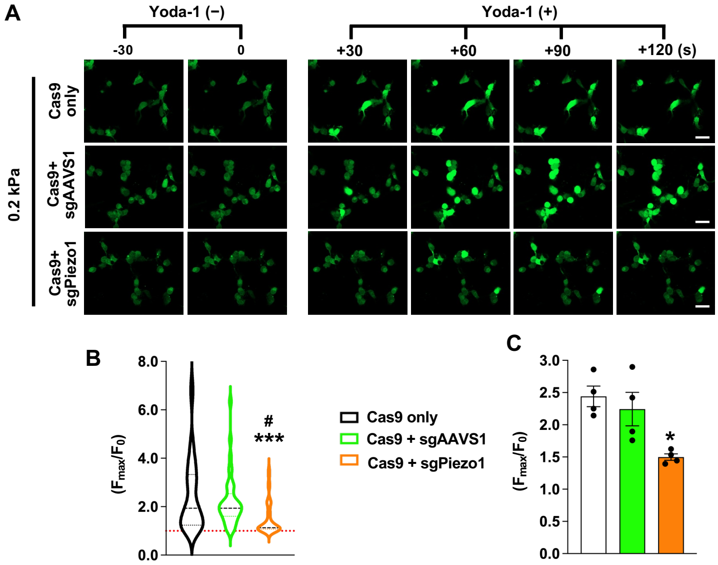


**Figure S12. Validation of functional Piezo1 knockdown by Crispr/Cas9 system**

**(A)** Representative time-lapse images of intracellular calcium uptake in NSCs cultured on 0.2 kPa hydrogel substrate. Piezo1 agonist Yoda-1 was added to the culture plate at 100 µM after 30 sec baseline recording. Calcium uptake was visualized using Fluo4-AM. Scale bars indicate 20 µm. **(B-C)** Quantitative graphs comparing the intracellular calcium levels in response to Piezo1 agonist stimulation. *** indicate $p < 0.001$ comparing Cas9 only with Cas9 + sgPiezo1, and # indicates $p < 0.05$ comparing Cas9 + sgScr with Cas9 + sgPiezo1 (B). * indicates $p < 0.05$ (C) by one-way ANOVA followed by Tukey's *post hoc* analysis. N = 4 biological replicates (40 cells per replicate), derived from two independent cultures.

**Table S1. sgRNAs targeting rat Piezo1 used in this study.**

| Target site | sgRNA # | Target sequence - PAM (5' to 3') |
|---|---|---|
| *rPiezo1* | Exon1-#1 | GAGCCGCACGTGCTGGGCGC-CGG |
| | Exon1-#2 | AGCCGCACGTGCTGGGCGCC-GGG |
| | Exon1-#3 | AGCCCGGCGCCCAGCACGTG-CGG |
| | Exon1-#4 | TGCTGGGCGCCGGGCTCTAC-TGG |
| | Exon1-#5 | GCTACCCTGCACACTCCTGG-CGG |
| | Exon2-#1 | GAGAGCATTGAAGCGTAACA-GGG |
| | Exon2-#2 | AGAGAGCATTGAAGCGTAAC-AGG |
| | Exon2-#3 | ACGCTTCAATGCTCTCTCGT-TGG |
| | Exon2-#4 | ATGCTGTGTCTTGAGGGGCC-TGG |
| | Exon3-#1 | GGCAGAGTAGTGCACGAAGC-AGG |
| | Exon3-#2 | CATTCCGTCACAGGTCACAC-AGG |
| | Exon3-#3 | CCTCAGCCTCCTCTTCCTGG-TGG |
| | Exon3-#4 | CATGGGCCACCAGGAAGAGG-AGG |

**Table S2. Representative targeted deep-sequencing reads (5 most frequent indel patterns) from rPiezo1-exon1-#3 RNP-treated samples. The PAM sequence is shown in green, and the sgRNA sequence is underlined**

| Indel | Local Sequence | Frequency (%) |
|---|---|---|
| WT | TCGCACCTCCCTTTAGCCACGGCGGCGCGACCCGGGGCGGTCCCCCGGCCGCGGGGCATGGAGCCGCACGTGCTGGGCGCCGGGCTCTAC | |
| -60 | TCGCACCTCCCTTTAGC------------------------------------------------------------CGCCGGGCTCTAC | 41.32 |
| -48 | TCGCACCTCCCTTTAGCCACGGCGGCGCGAC------------------------------------------------CCGGGCTCTAC | 18.42 |
| -55 | TCGCACCTCCCTTTAGCCACGGCGG-------------------------------------------------------CGGGCTCTAC | 15.50 |
| -38 | TCGCACCTCCCTTTAGCCACGGCGGCGCGACCCGGGGCGGTCCCC--------------------------------------TCTCTAC | 13.83 |
| -49 | TCGCACCTCCCTTTAGCCACGGCGGCGCGACCC-------------------------------------------------GGCTCTAC | 11.06 |

**Table S3 Primers used for sgRNA screening in this study.**

| Target site | Primer-F (5' to 3') | Primer-R (5' to 3') |
|---|---|---|
| *rPiezo1* sgRNA exon1-#1~#5 | AGTGCCATGGCGCGCGGTTC | GAATCTCCTCGCCTATGCTC |
| *rPiezo1* sgRNA exon2-#1~#4 | GAACCGGGGAGGTACTGCTC | GTAGCTTTCTGTAGGGTACC |
| *rPiezo1* sgRNA exon3-#1~#4 | TGGACAGCGCACGGTCTACC | GTCGCTCCTCGATCACTTAC |

**Table S4 Primers used for PCR in this study.**

| Target gene | Primer-F (5' to 3') | Primer-R (5' to 3') |
|---|---|---|
| 18S rRNA | CGGCTACCACATCCAAGGAA | TGCTGGCACCAGACTTGCCCTC |
| Piezo1 | CAATGCTCTCTCGTTGGTCT | CACGTTTGCCCAAAGGTT |
| TRPP2 | AATATGACCAGGACGGCGAC | TGGCCACTGTCTTCATCGTC |
| TRPC1 | AAGCTTTTCTTGCTGCCGTG | CGTTCCATAAGTTTCTGACAACCG |
| TRPA1 | GCAGCTTATTGCCTTCACAA | TTTGCGCAAGTACCAGAATG |
| TRPV4 | GGGAACCATCCACAGGGAAG | CTGTCGCCTCATATCGGCTT |

**Movie S1.**
Speed of gelation by injecting 10% and 16% I-5 hydrogels mixed with dye into a jar containing warm water at 37°C

**Movie S2.**
Spontaneous calcium oscillation of NSCs grown on plastic dish

**Movie S3.**
Spontaneous calcium oscillation of NSCs grown on 25 kPa and 0.2 kPa hydrogel substrate

**Movie S4.**
Spontaneous calcium oscillation of NSCs after cytoD treatment

**Movie S5.**
Calcium influx after Yoda1 treatment in NSCs grown on 0.2 kPa hydrogel substrate
